# Automated characterization of noise distributions in diffusion MRI data

**DOI:** 10.1101/686436

**Authors:** Samuel St-Jean, Alberto De Luca, Chantal M. W. Tax, Max A. Viergever, Alexander Leemans

**Affiliations:** Image Sciences Institute, Department of Radiology, University Medical Center Utrecht, Heidelberglaan 100, 3584 CX Utrecht, the Netherlands; Cardiff University Brain Research Imaging Centre (CUBRIC), School of Psychology, Cardiff University, Maindy Road, Cardiff, CF24 4HQ, United Kingdom

**Keywords:** Diffusion MRI, Noise estimation, Parallel acceleration, Gamma distribution, GRAPPA, SENSE

## Abstract

Knowledge of the noise distribution in magnitude diffusion MRI images is the centerpiece to quantify uncertainties arising from the acquisition process. The use of parallel imaging methods, the number of receiver coils and imaging filters applied by the scanner, amongst other factors, dictate the resulting signal distribution. Accurate estimation beyond textbook Rician or noncentral chi distributions often requires information about the acquisition process (e.g., coils sensitivity maps or reconstruction coefficients), which is usually not available. We introduce two new automated methods using the moments and maximum likelihood equations of the Gamma distribution to estimate noise distributions as they explicitly depend on the number of coils, making it possible to estimate all unknown parameters using only the magnitude data. A rejection step is used to make the framework automatic and robust to artifacts. Simulations using stationary and spatially varying noncentral chi noise distributions were created for two diffusion weightings with SENSE or GRAPPA reconstruction and 8, 12 or 32 receiver coils. Furthermore, MRI data of a water phantom with different combinations of parallel imaging were acquired on a 3T Philips scanner along with noise-only measurements. Finally, experiments on freely available datasets from a single subject acquired on a 3T GE scanner are used to assess reproducibility when limited information about the acquisition protocol is available. Additionally, we demonstrated the applicability of the proposed methods for a bias correction and denoising task on an in vivo dataset acquired on a 3T Siemens scanner. A generalized version of the bias correction framework for non integer degrees of freedom is also introduced. The proposed framework is compared with three other algorithms with datasets from three vendors, employing different reconstruction methods. Simulations showed that assuming a Rician distribution can lead to misestimation of the noise distribution in parallel imaging. Results on the acquired datasets showed that signal leakage in multiband can also lead to a misestimation of the noise distribution. Repeated acquisitions of in vivo datasets show that the estimated parameters are stable and have lower variability than compared methods. Results for the bias correction and denoising task show that the proposed methods reduce the appearance of noise at high b-value. The proposed algorithms herein can estimate both parameters of the noise distribution automatically, are robust to signal leakage artifacts and perform best when used on acquired noise maps.

## 1. Introduction

Diffusion magnetic resonance imaging (dMRI) is a non invasive imaging technique which allows probing microstructural properties of living tissues. Advances in parallel imaging techniques (Pruessmann et al., 1999; Griswold et al., 2002), such as accelerated acquisitions (e.g., partial k-space (Storey et al., 2007), multiband imaging (Nunes et al., 2006; Moeller et al., 2010) and compressed sensing (Lustig et al., 2007; Paquette et al., 2015)), have greatly reduced the inherently long scan time in dMRI. New acquisition methods and pulse sequences in dMRI are also pushing the limits of spatial resolution while reducing scan time (Holdsworth et al., 2019), which also affects the signal distribution in ways that are challenging to model. Estimation of signal distributions deviating from theoretical cases is challenging and oftentimes requires information such as coil sensitivities or reconstruction matrices. This information may not be recorded at acquisition time or is even not available from the scanner, making techniques relying on these parameters difficult to apply in practice. Even though the magnitude signal model is still valid nowadays, the use of image filters (Dietrich et al., 2008), acceleration methods subsampling k-space (e.g., the SENSE (SENsitivity ENcoding) (Pruessmann et al., 1999), GRAPPA (GeneRalized Autocalibrating Partial Parallel Acquisition) (Griswold et al., 2002; Heidemann et al., 2012) or the homodyne detection methods (Noll et al., 1991)) and spatial correlation between coil elements (Dietrich et al., 2008; Aja-Fernández et al., 2014) influence, amongst other factors, the parameters of the resulting signal distribution. See e.g., (Aja-Fernández and Tristán-Vega, 2015; Aja-Fernández et al., 2009) for a review on estimating noise distributions in MRI and common statistical distributions encountered therein.

With the recent trend towards open data sharing and large multicenter studies using standardized protocols (Duchesne et al., 2019; Emaus et al., 2015), differences in hardware, acquisition or reconstruction algorithms may inevitably lead to different signal distributions. This may affect large scale longitudinal studies investigating neurological changes due to these “scanner effects” (Sakaie et al., 2018) as the acquired data may be fundamentally different across sites in terms of statistical properties of the signal. Algorithms have been developed to mitigate these potential differences (Tax et al., 2019; Mirzaalian et al., 2018), but characterization of the signal distribution from various scanners is challenging due to the black box nature of the acquisition process, especially in routine clinical settings. While some recent algorithms for dMRI are developed to include information about the noise distribution (Collier et al., 2018; Sakaie and Lowe, 2017), there is no method, to the best of our knowledge, providing a fully automated way to characterize the noise distribution using information from the magnitude data itself only. Due to this gap between the physical acquisition process and noise estimation theory, noise distributions are either assumed as Rician (with parameter *σ_g_* related to the standard deviation) or noncentral chi (with fixed degrees of freedom *N*) and concentrate in estimating the noise standard deviation *σ_g_* (Veraart et al., 2016; Koay et al., 2009b; Tabelow et al., 2015). This assumption inevitably leads to misestimation of the true signal distribution as *N* and *σ_g_* are interdependent for some reconstruction algorithms (Aja-Fernández et al., 2013). Reconstruction filters preserving only the real part of the signal also cause *N* to deviate from the Rician noise distribution, producing instead a half-Gaussian signal distribution (Dietrich et al., 2008). Misestimation of the appropriate signal distribution could impact subsequent processing steps such as bias correction (Koay et al., 2009a), denoising (St-Jean et al., 2016) or diffusion model estimation (Zhang et al., 2012; Landman et al., 2007; Sakaie and Lowe, 2017), therefore negating potential gains in statistical power from analyzing datasets acquired in different centers or from different vendors.

In this work, we propose to estimate the parameters *σ_g_* and *N* from either the magnitude data or the acquired noise maps by using a change of variable to a Gamma distribution *Gamma*(*N*, 1) (Koay et al., 2009b), whose first moments and maximum likelihood equations directly depend on *N*. This makes the proposed method fast and easy to apply to existing data without additional information, while being robust to artifacts by rejecting outliers of the distribution. Preliminary results of this work have been presented at the annual meeting of the MICCAI (St-Jean et al., 2018b). This manuscript now contains additional theory, simulations including signal correlations and parallel acceleration, and experiments on phantoms and in vivo datasets acquired with parallel and multiband acceleration. As example applications, we perform bias correction and denoising on an in vivo dataset using the estimated distribution derived with each algorithm.

## 2. Theory

In this section, we introduce the necessary background on the Gamma distribution, its moments and maximum likelihood equations. Expressing the signal with a Gamma distribution highlights equations which can be solved to estimate parameters *σ_g_* and *N*.

### 2.1. Probability distribution functions of MRI data

To account for uncertainty in the acquisition process, the complex signal measured in k-space by the receiver coil array can be modeled with a separate additive zero mean Gaussian noise for each channel with identical variance 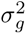 (Gudbjartsson and Patz, 1995). The signal acquired from the real and imaginary part of each coil in a reconstructed magnitude image can be expressed as (Constantinides et al., 1997)

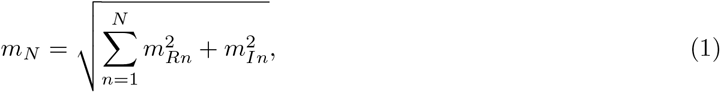

where *m_Rn_* and *m_In_* are the real and imaginary parts of the signal, respectively, as measured by coil number *n*, *N* is the number of degrees of freedom (which can be up to the number of coils in the absence of accelerated parallel imaging) and *m_N_* is the resulting reconstructed signal value for a given voxel. The magnitude signal can therefore be approximated by a noncentral chi distribution and has a probability density function (pdf) given by (Koay et al., 2009a; Dietrich et al., 2008)

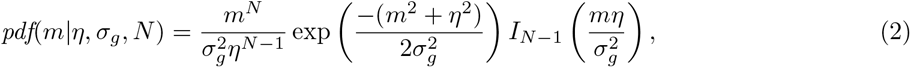

where *m* is the noisy signal value for a given voxel, *η* is the (unknown) noiseless signal value, *σ_g_* is the Gaussian noise standard deviation, *N* is the number of degrees of freedom and *I_ν_* (*z*) is the modified Bessel function of the first kind.

With the introduction of multiband imaging and other modern acquisition methods, parameters estimation of the magnitude data is not straightforward anymore. The number of degrees of freedom *N*, which is related to the number of receiver coils, likely deviates from heuristic estimation based on the actual number of coils as *N* also depends on the reconstruction technique employed (Sotiropoulos et al., 2013). The pdf of the magnitude data can be modeled by considering spatially varying degrees of freedom *N_eff_* and standard deviation *σ_eff_* (also called the *effective* values) and we generally have *N_eff_ ≤ N*, (Dietrich et al., 2008; Aja-Fernández et al., 2014).

The noncentral chi distribution includes the Rician (*N* = 1), the Rayleigh (*N* = 1*, η* = 0) and the central chi distribution (*η* = 0) as special cases (Dietrich et al., 2008). The pdf of the central chi distribution is given by

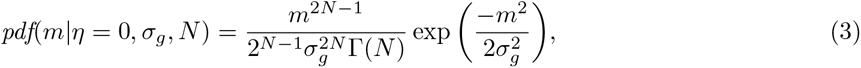

where Γ(*x*) is the Gamma function. With a change of variable introduced by (Koay et al., 2009b), Eq. (3) can be rewritten as a Gamma distribution *Gamma*(*N*, 1) with 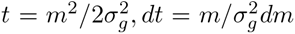 which has a pdf given by

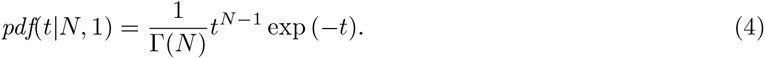

Eq. (4) only depends on *N*, which can be estimated from the sample values.

### 2.2. Parameter estimation using the method of moments and maximum likelihood The method of moments

The pdf of *Gamma*(*α, β*) is defined as

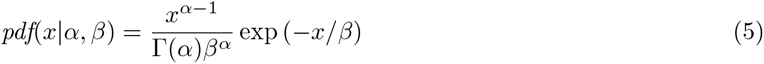

and has mean *µ_gamma_* and variance 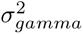 given by

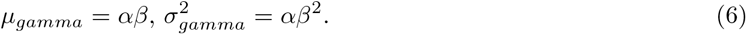

Another useful identity comes from the sum of Gamma distributions, which is also a Gamma distribution (Weisstein, 2017) such that if *t_i_* ~ *Gamma*(*α_i_, β*), then

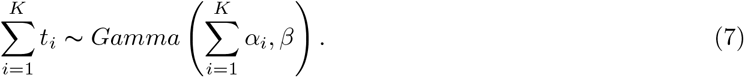

From Eq. (6), we obtain that the mean and the variance of the distribution *Gamma*(*N*, 1) are equal with a theoretical value of *N*. That is, we can estimate the Gaussian noise standard deviation *σ_g_* and the number of coils *N* from the sample moments of the magnitude images themselves, provided we can select voxels without any signal contribution i.e., where *η* = 0. Firstly, *σ_g_* can be estimated from Eq. (6) as

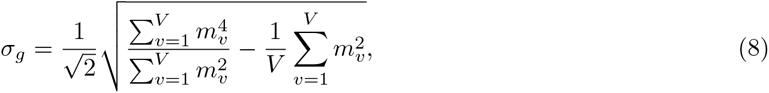

where *V* is the number of identified noise only voxels and *m_v_* the value of such a voxel, see Appendix A for the derivations. Once *σ_g_* is known, *N* can be estimated from the sample mean of those previously identified voxels as

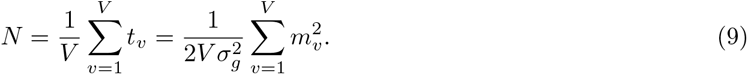

Derivations of Eqs. (8) and (9) are detailed in Appendix A.

#### Maximum likelihood equations for the Gamma distribution

Estimation based on the method of maximum likelihood yields two equations for estimating *α* and *β*. Rearranging the equations for a Gamma distribution will give Eq. (9) and a second implicit equation for *N* that is given by (Thom, 1958)

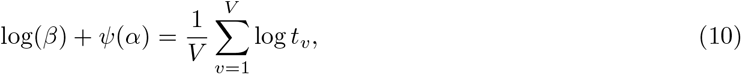

where 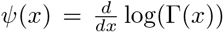 is the digamma function. For the special case *Gamma*(*N*, 1), we can rewrite Eq. (10) as

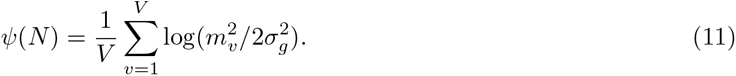

Combining Eq. (9) and Eq. (11), we also have an implicit equation to find *σ_g_*

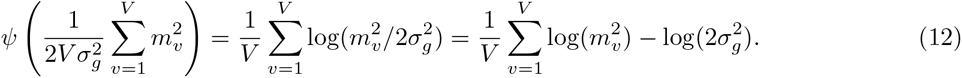

As Eqs. (11) and (12) have no closed form solution, they can be solved numerically e.g., using Newton’s method. See Appendix A for practical implementation details.

## 3. Material and Methods

### 3.1. Automated and robust background separation

The equations we presented in Section 2.1 are only valid when *η* = 0 by construction and assume that each selected voxel *m_v_* belongs to the same Gamma distribution. Following a methodology similar to (Koay et al., 2009b), we assume that each 2D slice with the same spatial location belongs to the same statistical distribution throughout each 3D volume. This practical assumption allows selecting a large number of noise only voxels for computing statistics as well as identifying (and subsequently discarding) potential slice acquisition artifacts which may affect one volume, but not the rest of the acquisition. Using Eq. (7), the sum of all diffusion weighted images (DWIs) can be used to separate the voxels belonging to the Gamma distribution *Gamma*(*KN*, 1), where *K* is the number of acquired DWIs, from the voxels not in that specific distribution with a rejection step using the inverse cumulative distribution function^1^(cdf). In the particular case *Gamma*(*KN*, 1) at a probability level *p*, the inverse cdf is *icdf*(*α, p*) = *P*^−1^(*α, p*), where *P*^−1^ is the inverse lower incomplete regularized gamma function^2^. This relationship can be used to identify potential outliers, such as voxels which contain non background signal, by excluding any voxel *m_v_* whose value does not fall between *λ*_−_ = *icdf*(*α, p*/2) and *λ*_+_ = *icdf*(*α*, 1 − *p/*2), i.e., *m_v_* is an outlier if *m*_*v*_ < *λ*_−_ or *m*_*v*_ > *λ*_+_.

To provide a better understanding of the change of variable 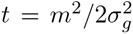, Fig. 1 shows the histogram for a synthetic dataset at b = 3000 s/mm^2^, which will be detailed later in Section 3.2. Voxels belonging to the background are easily separated in terms of the Gamma distribution after transformation, thus allowing estimation of parameters from voxels truly belonging to the noise distribution, see Appendix C and (St-Jean et al., 2018b) for technical details. Our implementation of the proposed algorithm is freely available^3^ (St-Jean et al., 2019).

**Figure 1:**
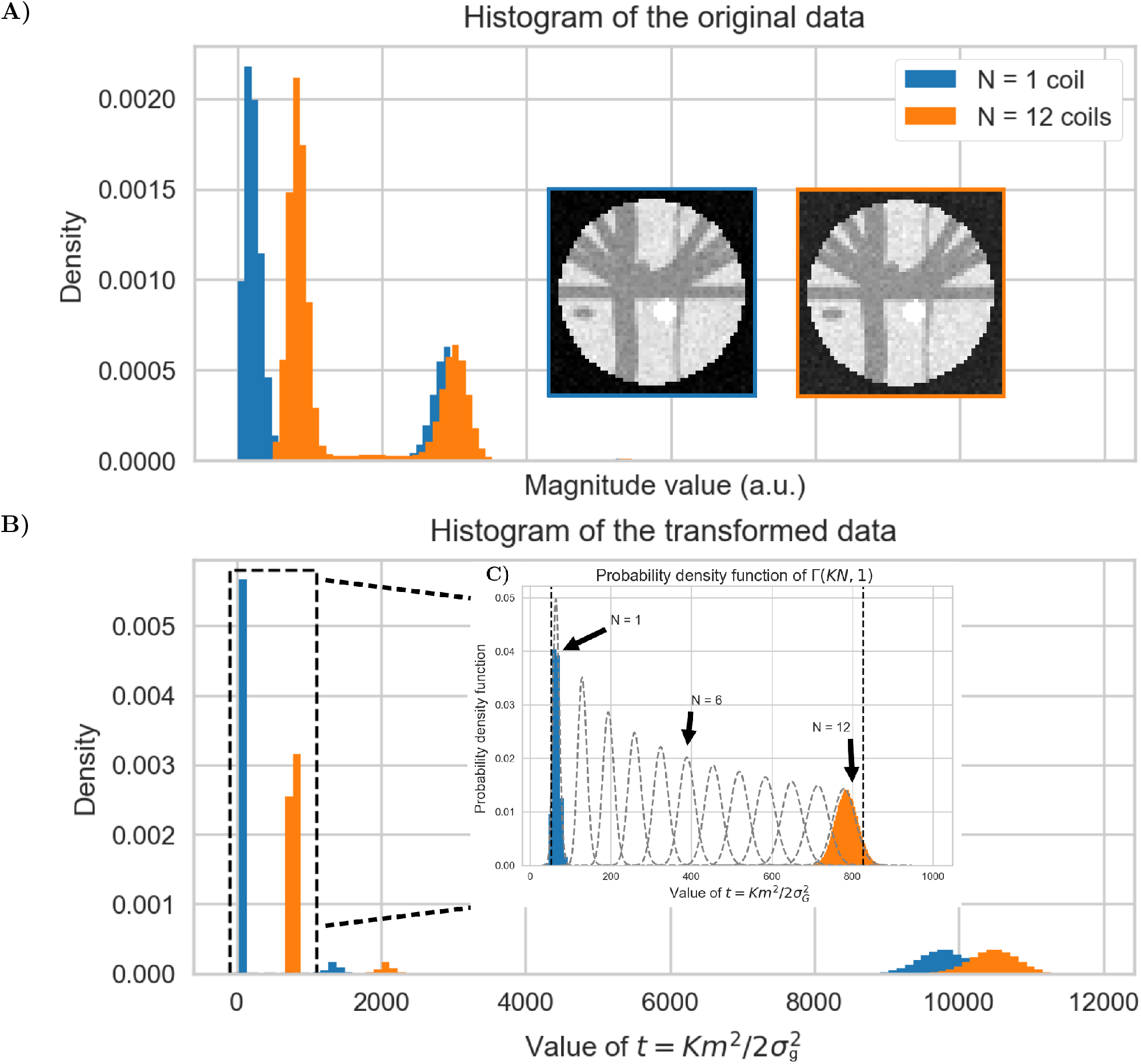
Histogram of the synthetic data at b = 3000 s/mm^2^ **A)** before the change of variable to a Gamma distribution and **B)** after the change of variable to a Gamma distribution for *N* = 1 and *N* = 12 with the true value of *σ*_*g*_. Summing all *K* DWIs together separates the background voxels from the rest of the data, which follows a Gamma distribution *Gamma*(*KN*, 1) by construction. In **C)**, a view of the left part from **B)** with the theoretical histograms of Gamma distributions from *N* = 1 up to *N* = 12. The black dotted lines represent the lower bound *λ*_−_ to the upper bound *λ*_+_, with *p* = 0.05*, N_min_* = 1 and *N_max_* = 12. This broad search covers the background voxels in both cases while excluding remaining voxels which do not belong to the distribution *Gamma*(*KN*, 1).

### 3.2. Datasets and experiments

#### Synthetic phantom datasets

Two synthetic phantom configurations from previous dMRI challenges were used. The first simulations were based on the ISBI 2013 HARDI challenge using phantomas (Caruyer et al., 2014). We used the given 64 gradient directions to generate two separate noiseless single-shell phantoms with either b = 1000 s/mm^2^ or b = 3000 s/mm^2^ and an additional b = 0 s/mm^2^ volume. The datasets were then corrupted with Rician (*N* = 1) and noncentral chi noise profiles (*N* = 4, 8 and 12), both stationary and spatially varying, at a signal-to-noise ratio (SNR) of 30 according to

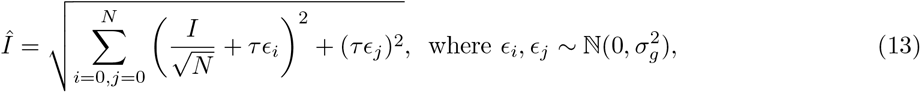

where *I* is the noiseless volume, 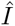 is the resulting noisy volume, *τ* is a mask for the spatial noise pattern, 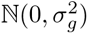 is a Gaussian distribution of mean 0 and variance 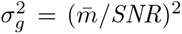 and 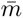 is the average signal value of the b = 0 s/mm^2^ image inside the white matter. In the stationary noise case, *τ* is set to 1 so that the noise is uniform. For the spatially varying noise case, *τ* is a sphere with a value of 1 in the center up to a value of 1.75 at the edges of the phantom, thus generating a stronger noise profile outside the phantom than for the stationary noise case. Since all datasets are generated at SNR 30, the noise standard deviation *σ_g_* is the same even though the b-value or number of coils *N* is different, but the magnitude standard deviation of the noise only voxels *σ_m_* is lower than *σ_g_*.

The second set of synthetic experiments is based on the ISMRM 2015 tractography challenge (Maier-Hein et al., 2017) which consists of 25 manually delineated white matter bundles. Ground truth data consisting of 30 gradient directions at either b = 1000 s/mm^2^ or b = 3000 s/mm^2^ and 3 b = 0 s/mm^2^ images at a resolution of 2 mm isotropic was generated using Fiberfox (Neher et al., 2014) without artifacts or subject motion. Subsequent noisy datasets were created at SNR 20 by simulating an acquisition with 8, 12 and 32 coils using the parallel MRI simulation toolbox^4^ with SENSE (Pruessmann et al., 1999) or GRAPPA (Griswold et al., 2002) reconstructions with an acceleration factor of *R* = 2. The SENSE simulated datasets also included spatial correlations between coils of *ρ* = 0.1, increasing the spatially varying effective noise standard deviation *σ_g_* and keeping the signal Rician distributed (*N* = 1). For the GRAPPA reconstructed datasets, 32 calibrating lines were sampled in the k-space center, neglecting spatial correlations (*ρ* = 0) as it is a k-space method (Aja-Fernández and Tristán-Vega, 2015). The resulting effective values of *N* and *σ_g_* will be both spatially varying. We additionally generated 33 synthetic noise maps per dataset by setting the underlying signal value to *η* = 0 and performing the reconstruction using the same parameters as the DWIs. All generated datasets are available online (St-Jean et al., 2018a).

#### Acquired phantom datasets

We acquired phantom images of a bottle of liquid on a 3T Philips Ingenia scanner using a 32 channels head coil with a gradient strength of 45 mT/m. We varied the SENSE factor from *R* = 1, 2 or 3 and multiband acceleration factors from no multiband (*MB*), *MB* = 2 or *MB* = 3 while fixing remaining acquisition parameters to investigate their influence on the resulting signal distributions, resulting in 9 different acquisitions. The datasets consist of 5 b = 0 s/mm^2^ volumes and 4 shells with 10 DWIs each at b = 500 s/mm^2^, b = 1000 s/mm^2^, b = 2000 s/mm^2^ and b = 3000 s/mm^2^ with a voxel size of 2 mm isotropic and TE / TR = 135 ms / 5000 ms, Δ*/δ* = 66.5 ms / 28.9 ms. Six noise maps were also acquired during each of the experiments by disabling the RF pulse and gradients of the sequence. The acquired phantom datasets are also available (St-Jean et al., 2018a).

#### In vivo datasets

A dataset consisting of four repetitions of a single subject^5^ was also used to assess the reproducibility of noise estimation without *a priori* knowledge (Poldrack et al., 2015). This is the dataset we previously used in our MICCAI manuscript (St-Jean et al., 2018b). The acquisition was performed on a GE MR750 3T scanner at Stanford university, where a 3x slice acceleration with blipped-CAIPI shift of FOV/3 was used, partial Fourier 5/8 with a homodyne reconstruction and a minimum TE of 81 ms. Two acquisitions were made in the anterior-posterior phase encode direction and the two others in the posterior-anterior direction. The voxelsize was 1.7 mm isotropic with 7 b = 0 s/mm^2^ images, 38 volumes at b = 1500 s/mm^2^ and 38 volumes at b = 3000 s/mm^2^. As the acquisition used a homodyne filter to fill the missing k-space, this should lead in practice to a half Gaussian noise profile, a special case of the noncentral chi distribution with *N* = 0.5, due to using only the real part of the signal for the final reconstruction (Chap. 13 Bernstein et al., 2004; Noll et al., 1991; Dietrich et al., 2008).

In addition, one dataset acquired on a 3T Siemens Connectom scanner from the 2017 MICCAI harmonization challenge^6^ consisting of 16 b = 0 s/mm^2^ volumes and 3 shells with 60 DWIs each at b = 1200 s/mm^2^, b = 3000 s/mm^2^ and b = 5000 s/mm^2^ was used (Tax et al., 2019). The voxel size was 1.2 mm isotropic with a pulsed-gradient spin-echo echo-planar imaging (PGSE-EPI) sequence and a gradient strength of 300 mT/m. Multiband acceleration *MB* = 2 was used with GRAPPA parallel imaging with *R* = 2 and an adaptive combine reconstruction employing a 32 channels head coil. Other imaging parameters were TE / TR = 68 ms / 5400 ms, Δ*/δ* = 31.1 ms / 8.5 ms, bandwidth of 1544 Hz/pixel and partial Fourier 6/8.

#### Noise estimation algorithms for comparison

To assess the performance of the proposed methods, we used three other noise estimation algorithms previously used in the context of dMRI. Default parameters were used for all of the algorithms as done in St-Jean et al. (2018b). The local adaptive noise estimation (LANE) algorithm (Tabelow et al., 2015) is designed for noncentral chi signal estimation, but requires *a priori* knowledge of *N*. Default parameters were used with *k*^*^ = 20 as recommended. Since the method works on a single 3D volume, we only use the b = 0 s/mm^2^ image for all of the experiments to limit computations as the authors concluded that the estimates from a single DWI are close to the mean estimate. We also use the Marchenko-Pastur (MP) distribution fitting on the principal component analysis (PCA) decomposition of the diffusion data, which is termed MPPCA (Veraart et al., 2016). In all experiments, we used the suggested default local window size of 5 × 5 × 5. Finally, we also compare to the Probabilistic Identification and Estimation of Noise (PIESNO) (Koay et al., 2009b), which originally proposed the change of variable to the Gamma distribution that is at the core of our proposed method. PIESNO requires knowledge of *N* (which is kept fixed by the algorithm) to iteratively estimate *σ_g_* until convergence by removing voxels which do not belong to the distribution *Gamma*(*N*, 1) for a given slice. We set *p* = 0.05 and *l* = 50 for the initial search of *σ_g_* in PIESNO and our proposed method, with additional parameters set to *N_min_* = 1 and *N_max_* = 12 for all cases. When estimating distributions from noise maps, we compute values in small local windows of size 3 × 3 × 3. The list of the software implementations and their version used in this manuscript is available in the supplementary materials. To the best of our knowledge, ours is the first method which estimates both *σ_g_* and *N* jointly without requiring any prior information about the reconstruction process of the MRI scanner. Because PIESNO and LANE both *require* knowledge of the value of *N*, we set the correct value of *N* for the spatially varying noise phantom experiments and assume a Rician distribution by setting *N* = 1 for the remaining experiments when *N* is unknown. We quantitatively assess the performance of each method on the synthetic datasets by measuring the standard deviation of the noise and the percentage error inside the phantom against the known value of *σ_g_*, computed for each voxel as

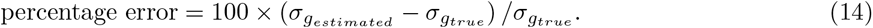

As PIESNO and our proposed methods estimate a single value per slice whereas MPPCA and LANE provide estimates from small spatial neighborhood, we report the mean value and the standard deviation of the error estimated inside the synthetic phantoms on each slice. For the acquired phantom datasets, we report the estimated noise distributions using both the DWIs and the measured noise maps for all 9 combinations of parallel imaging parameters that were acquired. For the in vivo datasets, we report once again the noise distributions estimated by each method. The reproducibility of the estimated distributions is assessed on the four GE datasets while the Connectom dataset is used to evaluate the performance of each compared algorithm on a bias correction and denoising task. In addition, we report *N* as estimated by our proposed methods for all cases.

#### Bias correction and denoising of the Connectom dataset

In a practical setting, small misestimation in the noise distribution (e.g., spatially varying distribution vs nature of the distribution) might not impact much the application of choice. We evaluate this effect of misestimation on the Connectom dataset with a bias correction and a denoising task. Specifically, we apply noncentral chi bias correction (Koay et al., 2009a) on the in vivo dataset from the CDMRI challenge using Eq. (B.4). The algorithm is initialized with a spherical harmonics decomposition of order 6 (Descoteaux et al., 2007) as done in (St-Jean et al., 2016). The data is then denoised using the non local spatial and angular matching (NLSAM) algorithm with 5 angular neighbors where each b-value is treated separately (St-Jean et al., 2016). Default parameters of a spatial patch size of 3 3 3 were used and the estimation of *σ_g_* as computed by each method was given to the NLSAM algorithm. For MPPCA, LANE and PIESNO, a default value of *N* = 1 was used and the value of *N* as computed by the moments and maximum likelihood equations for the proposed methods. The bias correction algorithm was also generalized for non integer values of *N* as detailed in Appendix B.

## 4. Results

We show here results obtained on the phantoms and in vivo datasets. The first set of simulations uses a sum of square reconstruction with stationary and spatially varying noise profiles. The second set of simulations includes SENSE and GRAPPA reconstructions, resulting in both spatially varying signal distribution profiles. Finally, the distributions estimated by each algorithm for the in vivo dataset are used for a bias correction and denoising task.

### 4.1. Synthetic phantom datasets

#### Simulations with a sum of squares reconstruction

Fig. 2 shows results from simulations with stationary and spatially varying noise profiles for all datasets as estimated inside the ISBI 2013 challenge phantom. For stationary noise profiles with *N* unknown, estimation of *σ_g_* is the most accurate for the proposed methods with an error of about 1%, followed by MPPCA making an error of approximately 5% and LANE of 15%. The error of PIESNO increases with the value of *N*, presumably due to misspecification in the signal distribution, whereas MPPCA and LANE are both stable in their estimation with increasing values of *N*. The proposed methods using equations based on the moments and maximum likelihood recovers the correct value of *σ_g_* in all cases with the lowest variance across slices, indicating that the estimated value of *σ_g_* is similar in all slices as expected. The same behavior is observed for PIESNO when *N* = 1, but the estimated *σ_g_* is larger than the correct value by two to three times when *N* is misspecified. In the spatially varying noise case where *N* is known, the moments, maximum likelihood equations and PIESNO all perform similarly with approximately 2% of error. LANE generally outperforms MPPCA except for the *N* = 12 case, but still misestimates *σ_g_* by approximately 15% and 25% respectively. Only the proposed methods and MPPCA are independent of correctly specifying *N*. Finally, Fig. 3 shows the estimated values of *N* by the proposed methods. Estimation generally follows the correct value, regardless of misestimation of *σ_g_*.

**Figure 2:**
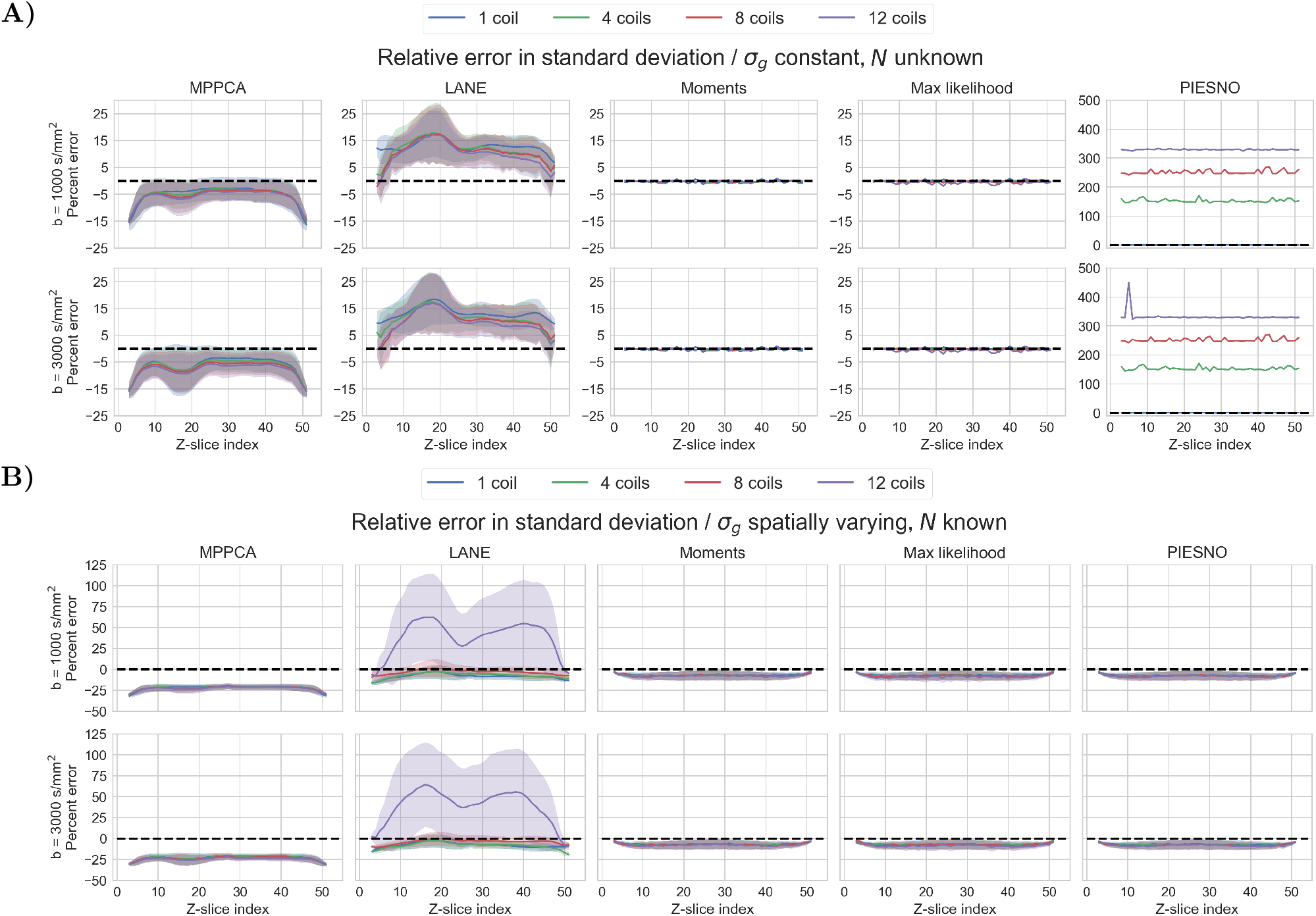
Percentage of error when the real value of *N* is unknown and *σ*_*g*_ is constant (in **A)**) and *N* is known with *σ*_*g*_ spatially varying (in **B)**) with the mean (solid line) and standard deviation (shaded area). All methods underestimate spatially varying *σ*_*g*_, except for LANE with *N* = 12 which overestimates it instead. On average, all methods are tied at around 5% of error with MPPCA reaching approximately 25% of error. Of interesting note, the proposed methods are tied with PIESNO when the correct value of *N* is given to the latter, but do not require an estimate of *N*, which is now an output instead of a prerequisite.

**Figure 3:**
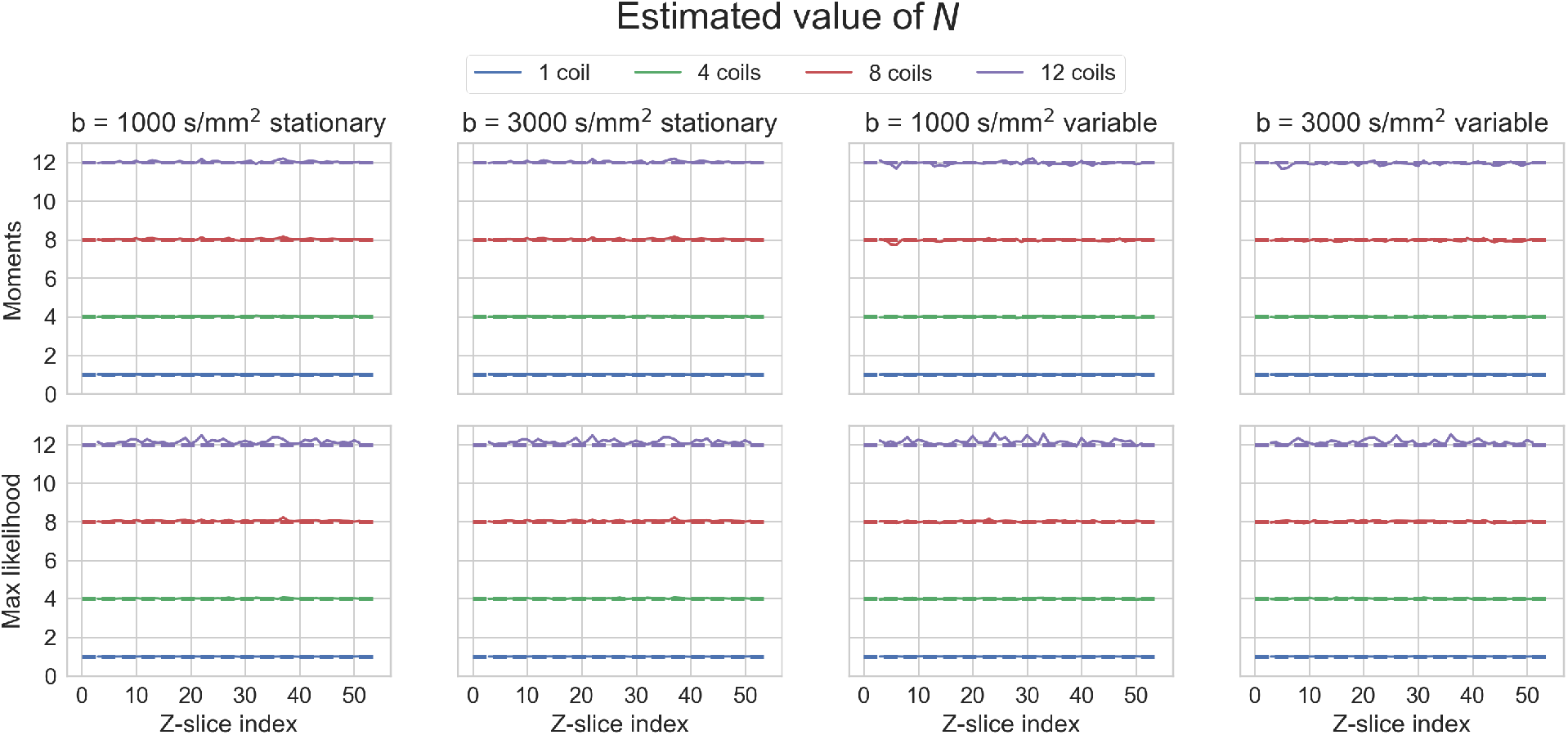
Estimated value of *N* using equations from the moments (top) and with maximum likelihood (bottom) for the proposed methods. Even for the spatially variable case where *σ*_*g*_ is slightly underestimated, the estimated values of *N* are stable and correspond to the real values used in the synthetic simulations in every case.

#### Simulations with parallel imaging

Fig. 4 shows the estimated values of *σ_g_* from a SENSE reconstruction and Fig. 5 shows the results for the GRAPPA reconstructed datasets. For SENSE, estimation using noise maps is the most precise for both proposed methods and PIESNO where the average error is around 0, followed by LANE when using DWIs as the input which results in 10% of overestimation. MPPCA generally underestimates *σ_g_* by around 15% for data at b = 1000 s/mm^2^ and 30% for data at b = 3000 s/mm^2^. LANE instead overestimates when using DWIs and underestimates *σ_g_* when using noise maps and knowing the correct value of *N* = 1. The proposed methods (the moments and maximum likelihood equations) and PIESNO are performing similarly, but PIESNO requires knowledge of *N* = 1. Estimation is also more precise for the three methods using the Gamma distribution (moments, maximum likelihood and PIESNO) than those using local estimations (MPPCA and LANE) and closest to the true values when using noise maps. In the case of GRAPPA, results are similar to the SENSE experiments with the exception of MPPCA being more precise than the compared methods for the b = 1000 s/mm^2^ case and performs equally well at b = 3000 s/mm^2^ as the proposed methods with an average error of about 20%. Results using LANE are similar with increasing number of coils when assuming *N* = 1, while the estimated value from PIESNO also increases with the number of coils as previously seen in Fig. 2. In this case, LANE overestimates *σ_g_* by around 50% when using DWIs, but performs similarly to MPPCA when estimating *σ_g_* from the noise maps. Estimation from noise maps using the moments or maximum likelihood equations is the most precise in all cases. The error of PIESNO increases with *N* as seen in Fig. 5 panel **C)**. This is caused by mistakenly including gray matter voxels of low intensity in the estimated distribution while they are correctly excluded automatically by the proposed methods. Finally, Fig. 6 shows the estimated values of *N_eff_* using the datasets from Figs. 4 and 5 by the proposed methods. For the SENSE case, the true value is a constant *N* = 1 by construction and the estimated values by both algorithms are on average correct with the maximum likelihood equations having the lowest variance. In the case of GRAPPA, values of *N* vary spatially inside the phantom and depend on the per voxel signal intensity, just as *σ_g_* does in Fig. 5. This leads to some overestimation when only background voxels are considered, with the best estimation obtained when using the noise maps. For simulations using 8 and 12 coils, estimated values of *N* are, in general, following the expected values. However, the spatially varying pattern can not be fully recovered as the correct value of *N* depends on the true signal intensity *η* in each voxel, which is not present when collecting noise only measurements.

**Figure 4:**
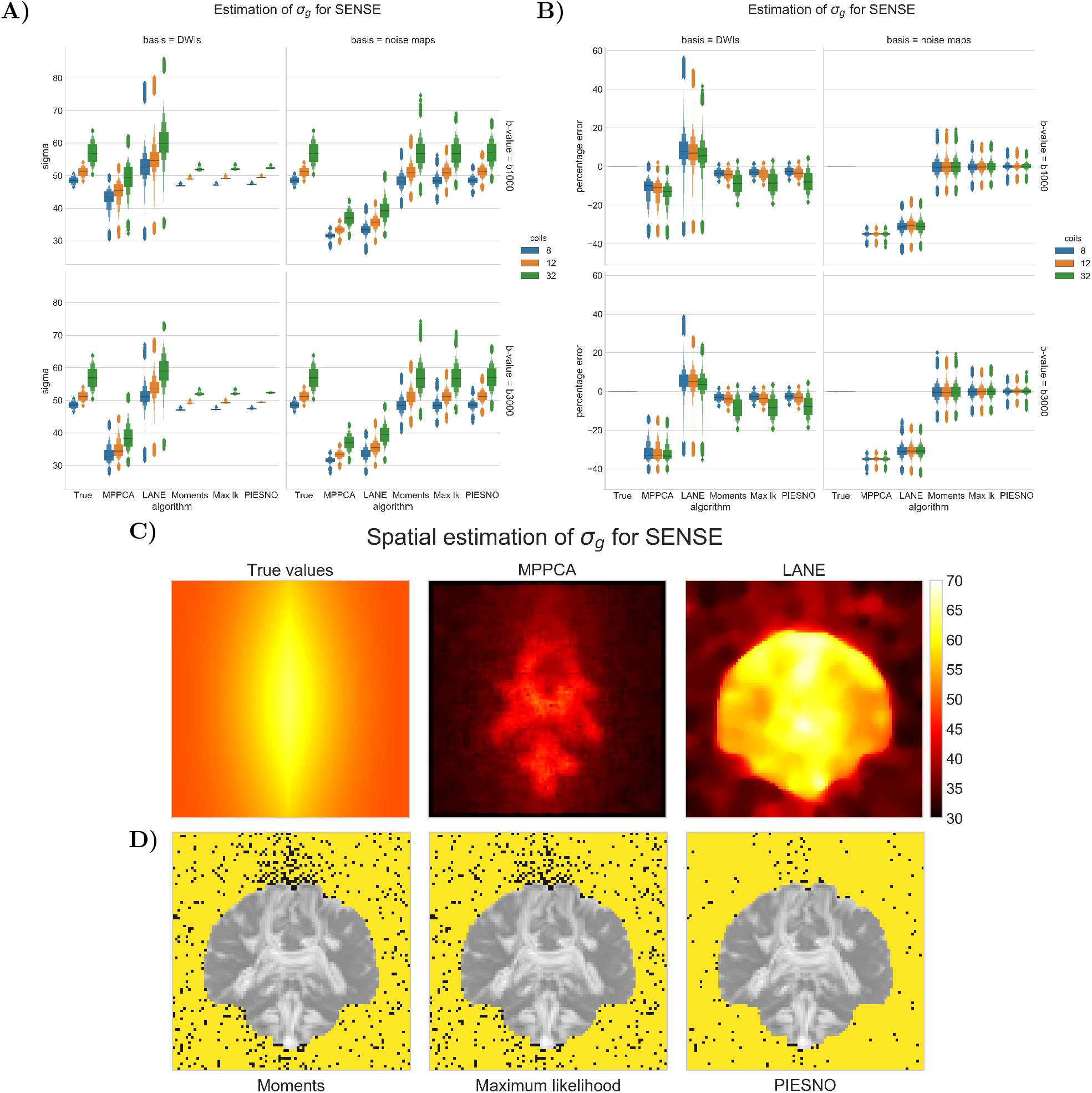
Estimation of the noise standard deviation *σ*_*g*_ (in **A**) and the percentage error (in **B**) inside the phantom only for each method using a SENSE reconstruction with 8, 12 or 32 coils. The left columns (basis = DWIs) shows estimation using all of the DWIs, while the right column (basis = noise maps) shows the estimated values from synthetic noise maps in small windows of size 3 × 3 × 3. Results for b = 1000 s/mm^2^ are on the top row, while the bottom row shows results for the b = 3000 s/mm^2^ datasets. Figure **C)** shows the spatially estimated values of *σ*_*g*_ using the b = 3000 s/mm^2^ dataset with 32 coils for a single slice from the true distribution and local estimation as done by MPPCA and LANE. The general trend shows that even though MPPCA and LANE misestimate *σ*_*g*_, they still follow the spatially varying pattern (lower at edges with the highest intensity near the middle) from the correct values. In **D)**, voxels identified as belonging to the same distribution Gamma(*N*, 1) are overlaid in yellow over the sum of all DWIs. Note how voxels containing signal from the DWIs are excluded by all three methods.

**Figure 5:**
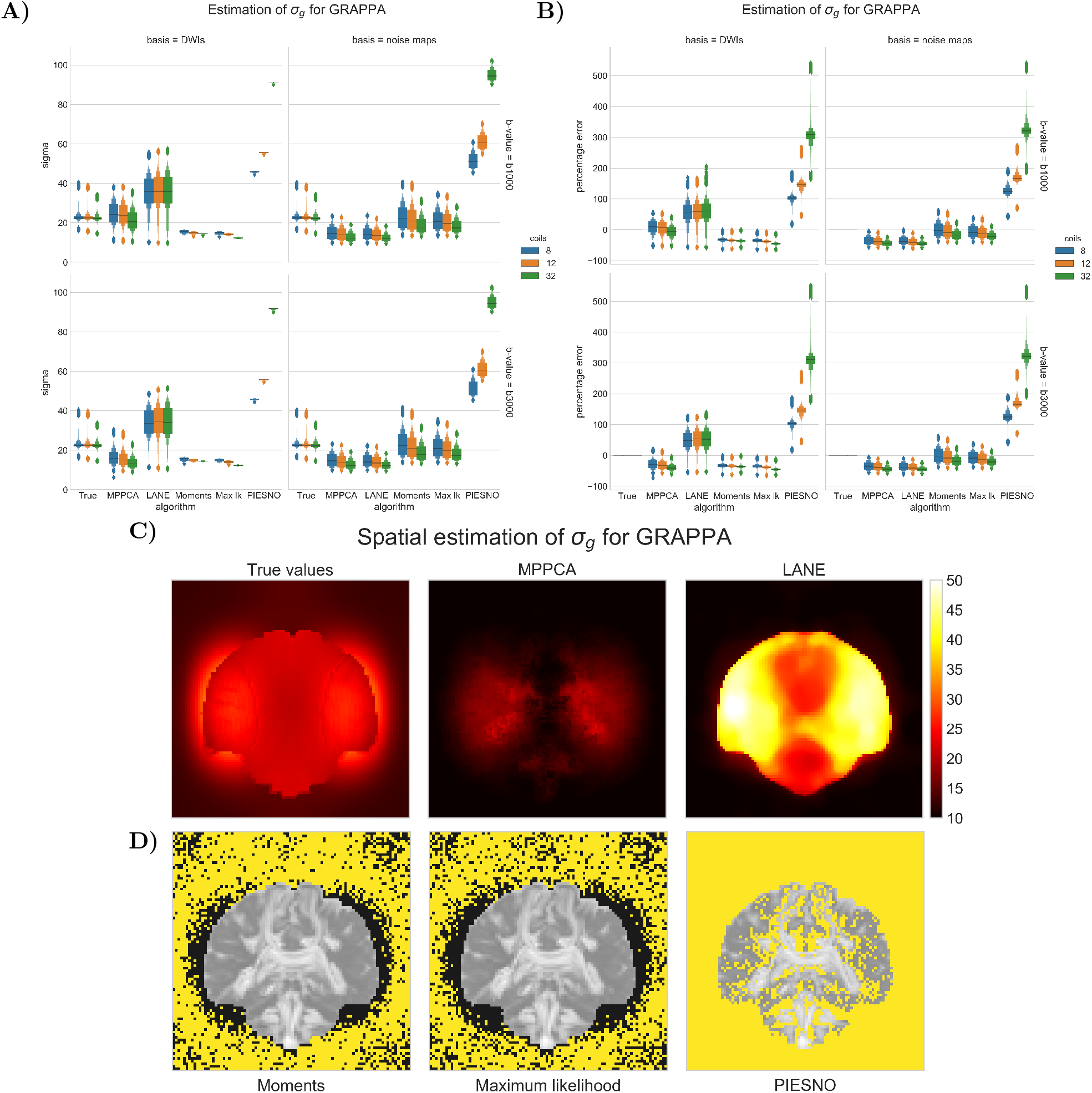
Estimation of the noise standard deviation *σ*_*g*_ (in **A**) and the percentage error (in **B**) inside the phantom only for each method using a GRAPPA reconstruction with 8, 12 or 32 coils, using the same conventions as Fig. 4. Figure **C)** shows the the true value of *σ*_*g*_ and the spatially estimated *σ*_*g*_ from MPPCA and LANE using the b = 3000 s/mm^2^ dataset with 32 coils for a single slice. There is once again a misestimation for both methods while following the correct spatially varying pattern. In **D)**, voxels identified as belonging to the same distribution Gamma(*N*, 1) are overlaid in yellow over the sum of all DWIs. Note how PIESNO mistakenly selects some low intensity voxels belonging to the gray matter, in addition to all of the voxels in the background, which causes an overestimation of *σ*_*g*_ with a fixed value of *N* = 1. Both proposed methods instead select voxels with small variations in intensity as belonging to the same distribution without mistakenly selecting gray matter voxels.

**Figure 6:**
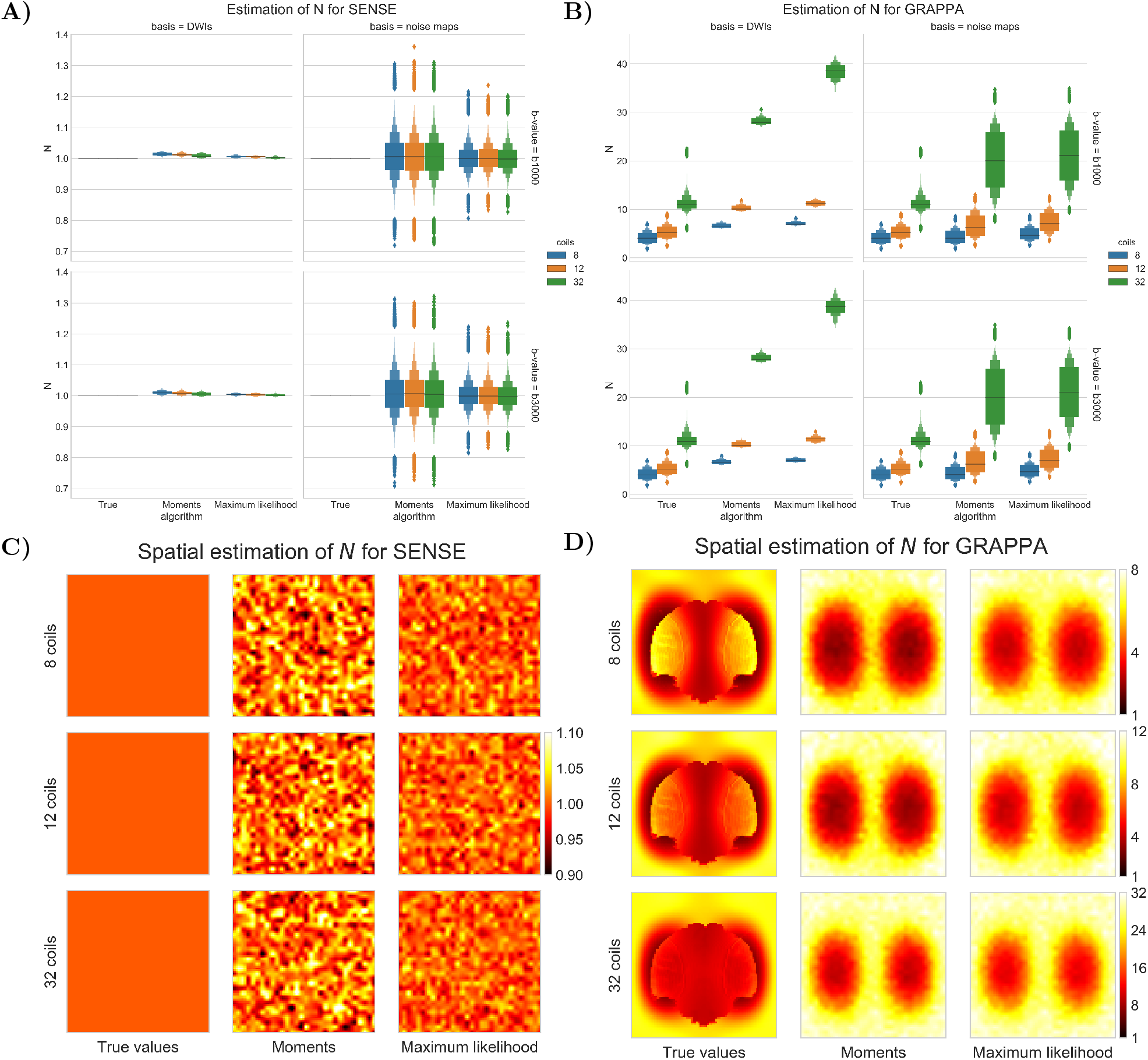
The estimated values of *N* for SENSE (left column) and GRAPPA (right column) for the b = 1000 s/mm^2^ (first row of boxplots) and b = 3000 s/mm^2^ (second row of boxplots) datasets. The left column shows results computed from the automatically selected background voxels (basis = DWIs), while the right column shows local estimation using noise maps (basis = noise maps). In **A)** and **B)**, the boxplot of *N* inside the phantom for the SENSE/GRAPPA algorithm with a spatial map of *N* shown in **C)** and **D)** computed using the noise maps from the b = 3000 s/mm^2^ datasets. Note how the colorbar is the same in **C)**, while each row of **D)** shares the same colorbar.

### 4.2. Acquired phantom datasets

Fig. 7 shows the estimated values of *σ_g_* for all methods with a SENSE acceleration of rate *R* = 1, 2 and 3 with multiband imaging at acceleration factors of *MB* = 2, *MB* = 3 or deactivated in panel **A)**. Results show that *σ_g_* increases with *R* and is higher when *MB* = 3 for *R* fixed, even if in theory *σ_g_* should be similar for a given *R* and increasing *MB*. Panel **C)** shows the estimated values of *σ_g_* when using noise maps as the input for *R* = 3 and *MB* = 3. As in the synthetic experiments, MPPCA and LANE have the lowest estimates for *σ_g_* with PIESNO and the proposed methods estimating higher values. Since the correct value is unknown, a reference sample slice of a noise map is also shown. When compared to values from the measured noise map, estimated values of *σ_g_* are approximately fivefold lower for MPPCA, four times lower for LANE and around half for the other methods. Estimation on the noise maps yields a value of around *N* = 1 for both proposed methods as seen in panels **B)** and **E)**, irrespective of the acceleration used. In the case of estimation using the DWIs, the range of estimated values is larger and increases at acceleration factors of *MB* = 2 and *R* = 2 or 3.

**Figure 7:**
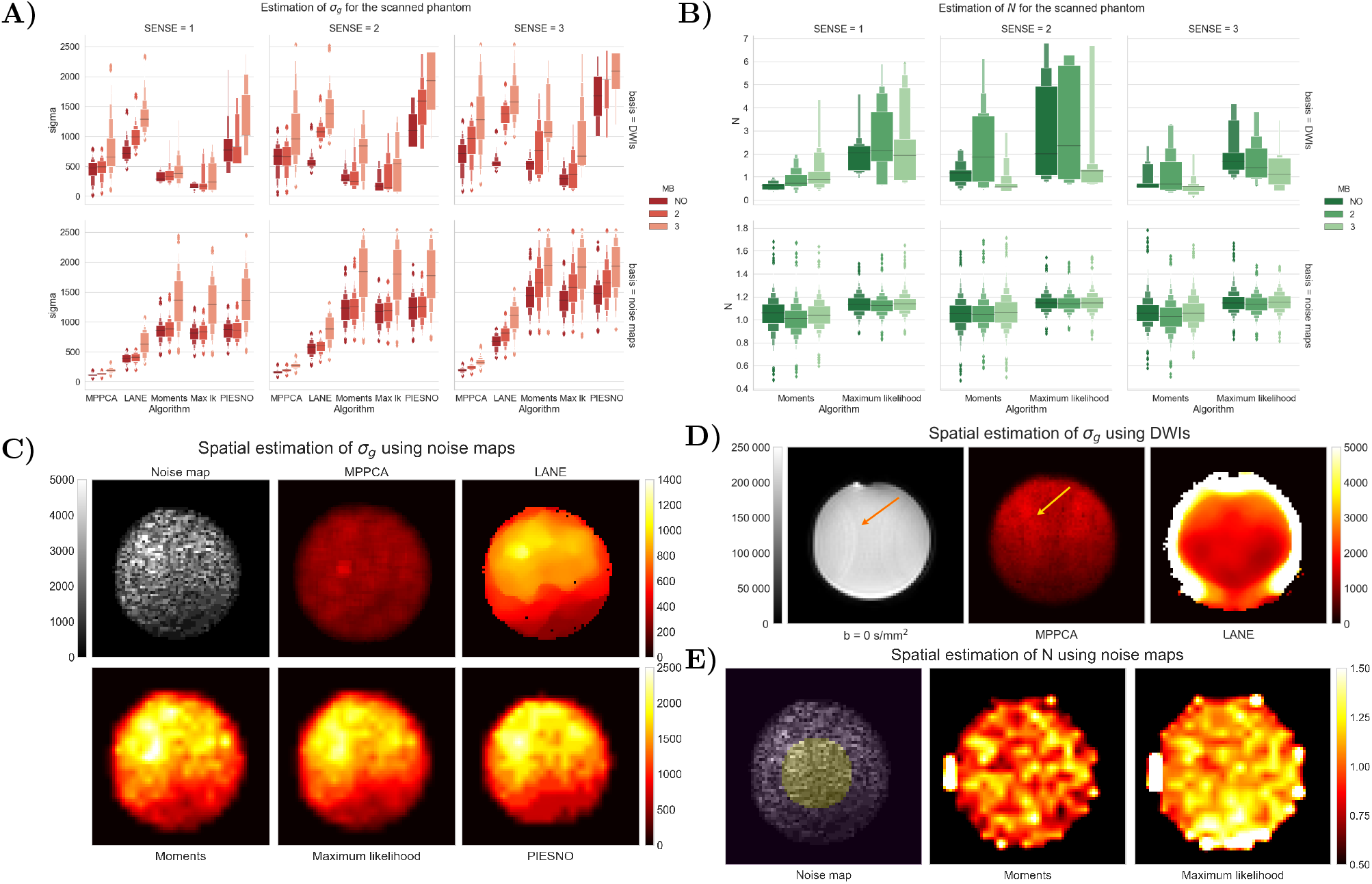
Estimation of noise distributions for the scanned phantom datasets inside a small ROI. Large outliers above the 95th percentile were removed to not skew the presented boxplots. In **A)**, the estimation of the noise standard deviation *σ*_*g*_ for each method using DWIs (top row) and using noise maps (bottom row). Each column shows an increasing SENSE factor, where *σ*_*g*_ increases (according to theory) with the square root of the SENSE factor. The different hues show an increasing multiband factor, which should not influence the estimation of *σ*_*g*_. For the case *MB* = 3, there may be signal leakage to adjacent slices, which would increase the measured values of *σ*_*g*_ even when the estimation uses only noise maps. In **B)**, boxplots for the values of *N* estimated by both proposed methodologies for the experiments shown in **A)**. Estimated values using noise maps are always close to 1 on average while estimations using DWIs seems to be affected by the possible signal leakage inherent to the use of multiband imaging. In **C)**, an axial slice of a noise map and estimated values of *σ*_*g*_ by all methods for the case *R* = 3 and *MB* = 3, which is the highest rate of acceleration from all of the investigated cases. Note the different scaling between the top and bottom row as MPPCA and LANE estimates of *σ*_*g*_ are two to three times lower than other methods. In **D)**, a b = 0 s/mm^2^ image of the phantom and spatially estimated values of *σ*_*g*_ for MPPCA and LANE. Note how some signal leakage (orange arrows) is affecting the b = 0 s/mm^2^ volume due to using *MB* = 3. In **E)**, location of the spherical ROI used for the boxplots overlaid on a noise map and spatially estimated values of *N* for both proposed methods. As less voxels are available near the borders of the phantom, estimating the noise distributions parameters results in lower precision.

### 4.3. In vivo datasets

#### Multiple datasets from a single subject

Fig. 8 shows the estimated value of *σ_g_* on four repetitions of the GE datasets for each method as computed inside a brain mask. The values from a b = 3000 s/mm^2^ volume (including background) is also shown as a reference for the values present at the highest diffusion weighting in the dataset. All methods show good reproducibility as their estimates are stable across the data. The value of *N* as computed by our proposed methods is also similar for all datasets with the median at *N* = 0.45 for the moments and *N* = 0.49 for the maximum likelihood equations. This corresponds to a half Gaussian distribution as would be obtained by a real part magnitude reconstruction (Dietrich et al., 2008). However, LANE recovered the highest values of *σ_g_* amongst all methods with a large variance and a median higher than the b = 3000 s/mm^2^ values, which might indicate overestimation in some areas. The median of MPPCA and the proposed methods are similar, while PIESNO estimates of *σ_g_* are approximately two times lower. This could indicate that specifying *N* = 1 was incorrect for these datasets, as PIESNO identified about 10 noise only voxels.

**Figure 8:**
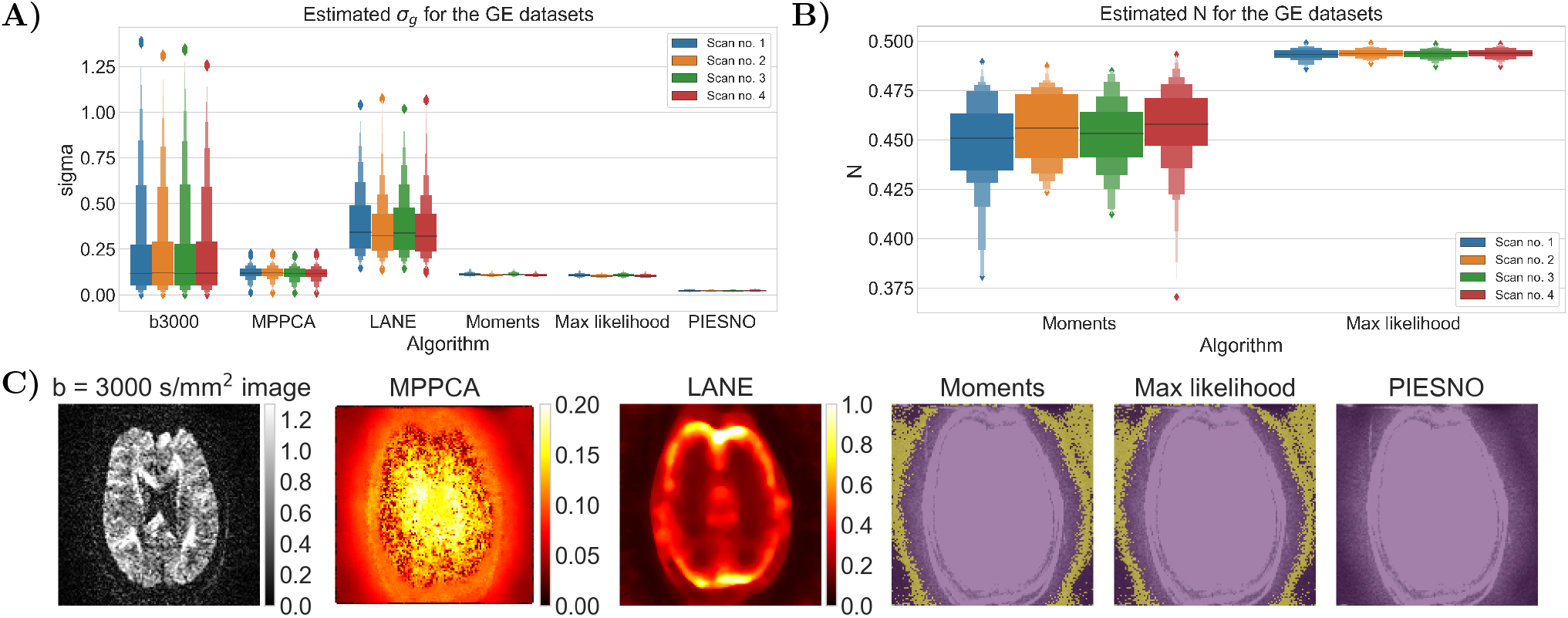
Estimation of the noise profiles on four repetitions of a single subject from a GE scanner. In **A)**, the baseline signal values of a b = 3000 s/mm^2^ volume and estimated values of *σ*_*g*_ for all methods inside a brain mask and **B)** estimated values of *N* by the proposed methods are shown. Note that the values for LANE and the b = 3000 s/mm^2^ volume were truncated at the 99 percentile to remove extreme outliers. In **C)**, an axial slice of a b = 3000 s/mm^2^ image from one dataset and the estimated values of *σ*_*g*_ for MPPCA and LANE. For the proposed methods and PIESNO, a mask of the identified background voxels (in yellow) overlaid on the data.

Fig. 9 shows an axial slice around the cerebellum and the top of the head which are corrupted by acquisition artifacts likely due to parallel imaging. Voxels containing artifacts were automatically discarded by both methods, preventing misestimation of *σ_g_* and *N*. The values computed from these voxels also offer a better qualitative fit than assuming a Rayleigh distribution or selecting non-brain data. We also timed each method to estimate *σ_g_* on one of the GE datasets using a standard desktop computer with a 3.5 GHz Intel Xeon processor. The runtime to estimate *σ_g_* (and *N*) was around 5 seconds for the maximum likelihood equations, 9 seconds for the moments equations, 11 seconds for PIESNO, 3 minutes for MPPCA and 18 minutes for LANE.

**Figure 9:**
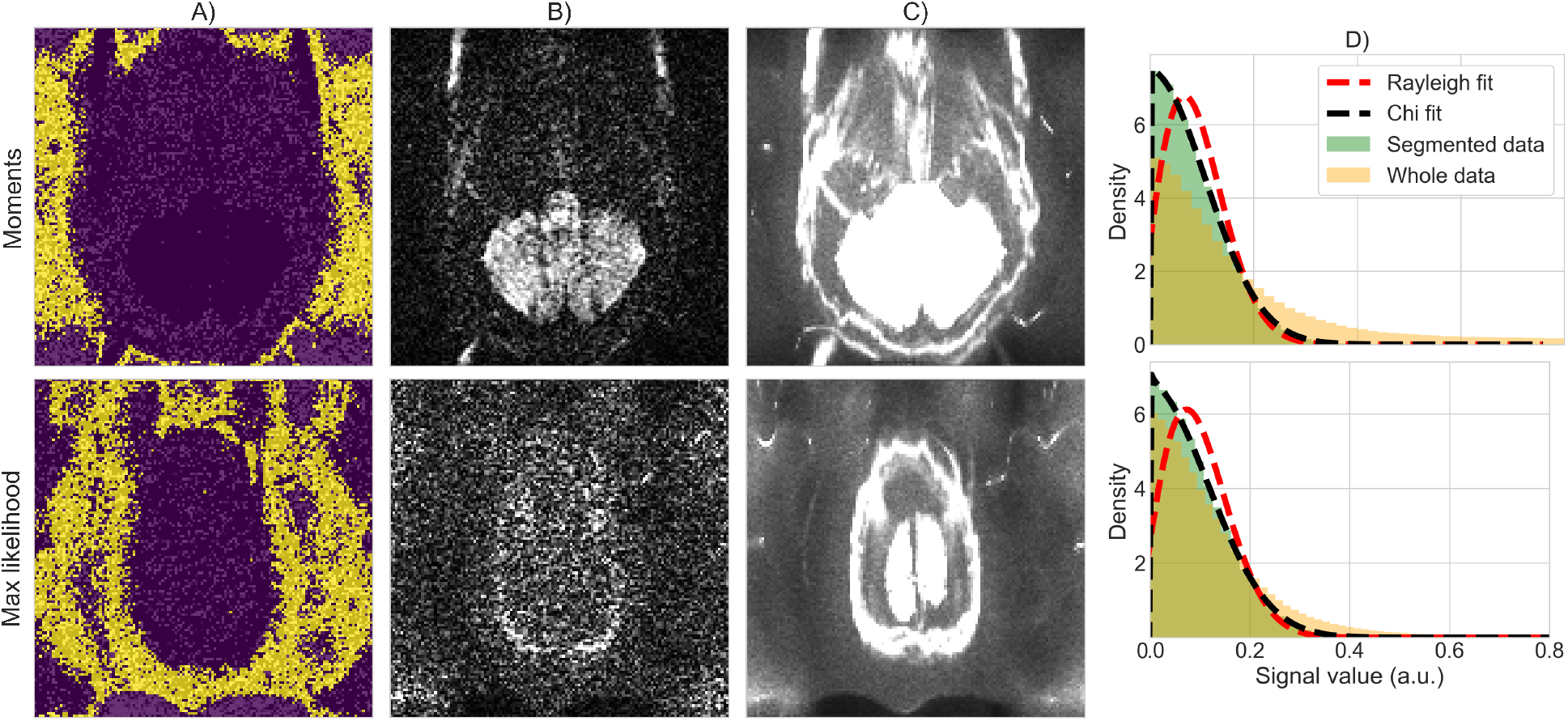
An axial slice in the cerebellum from one of the GE datasets. Voxels identified in **A)** as noise only (yellow) are free of artifacts in a single slice in **B)** or along the sum of all volumes in **C)**. In **D)**, the normalized density histogram using the selected voxels from **A)** (green) fit well a chi distribution (black dashed lines), while assuming a Rayleigh distribution (red dashed lines) or using all non brain voxels (orange) leads to a worse visual fit.

#### Estimation with a Connectom dataset

Fig. 10 shows in **A)** the estimated values of *σ_g_* inside a brain mask and in **B)** the values of *N* computed by the proposed methods. Estimated values of *σ_g_* vary by an order of magnitude between the different methods. In the case of MPPCA and LANE, the median of the estimates is higher than the reference b = 5000 s/mm^2^ data, while PIESNO and the proposed methods estimate values lower than the reference and have lower variability in their estimated values. For the estimation of *N*, recovered values are distributed close to 1 as is expected from an adaptive combine reconstruction providing a Rician distribution. Values estimated with the maximum likelihood equations have a lower variability than with the moments equations. In **C)**, the top row shows the b = 5000 s/mm^2^ volume and spatial maps of *σ_g_* as estimated by MPPCA and LANE. The bottom row shows voxels identified as pure noise (in light purple) using the moments, the maximum likelihood equations and PIESNO. Ghosting artifacts are excluded, but presumably affect estimation using the entire set of DWIs shown in the top row. Fig. 11 shows in **A)** the signal intensity after applying bias correction (left column) and after denoising (right column) for each volume ordered by increasing b-value. The top row (resp. bottom row) shows the mean (resp. standard deviation) as computed inside a white and gray matter mask. The mean signal decays with increasing b-value as expected, but the standard deviation of the signal does not follow the same trend in the cases of LANE. After denoising, the mean signal and its standard deviation decays once again as for the original data. Panel **B)** shows the average DWI at a given b-value for the original dataset and after denoising using the noise distribution from each method. Results are similar for all methods for the b = 0 s/mm^2^ datasets, but the overestimation of *σ_g_* by LANE produces missing values in the gray matter for b = 3000 s/mm^2^ and b = 5000 s/mm^2^. In general, averaging reduces the noise present at b = 0 s/mm^2^ and b = 1200 s/mm^2^ while only denoising is effective at b = 3000 s/mm^2^ and b = 5000 s/mm^2^. At b = 5000 s/mm^2^, the MPPCA denoised volume is of lower intensity than when obtained by the moments, maximum likelihood equations or PIESNO. This is presumably due to LANE and MPPCA estimating higher values of *σ_g_* than the three other methods. Finally, panel **C)** shows the absolute difference between the original and the denoised dataset obtained by each method. At b = 5000 s/mm^2^, LANE removes most of the signal in the gray matter mistakenly due to overestimating *σ_g_*. Other methods perform comparably well on the end result, despite estimates of *σ_g_* of different magnitude.

**Figure 10:**
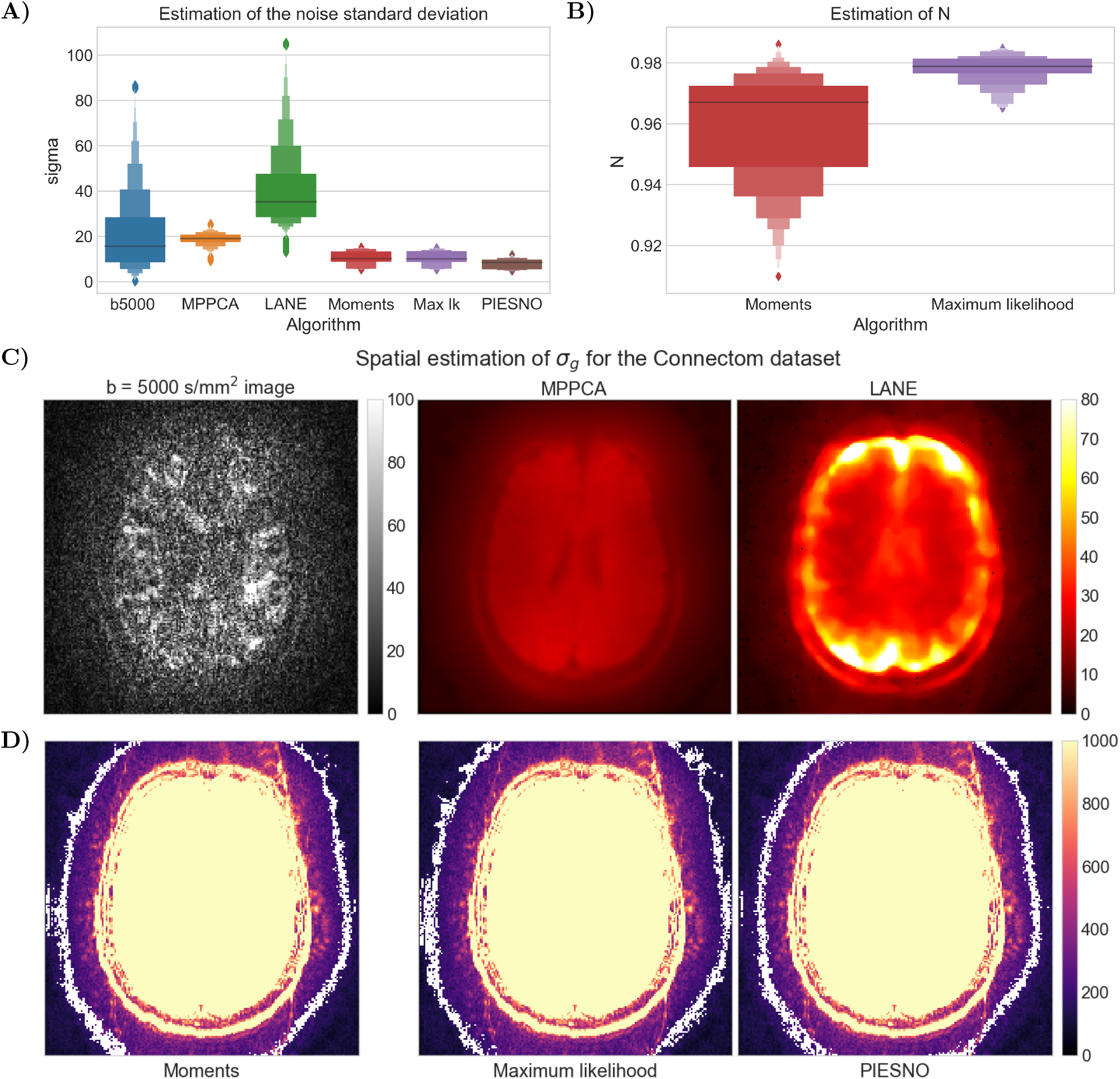
Estimation of noise distributions for the Connectom dataset. In **A)**, signal distribution of the original data and noise standard deviation *σ*_*g*_ for all methods, where data above the 99th percentile for the b = 5000 s/mm^2^ volume and LANE were discarded. In **B)**, values of *N* as estimated using the moments (in red) and by maximum likelihood (in purple). In **C)** on the top row, a b = 5000 s/mm^2^ volume and spatial estimation of *σ*_*g*_ as measured by MPPCA and LANE. In **D)**, voxels identified as containing only noise (in white) by the moments, maximum likelihood and PIESNO overlaid on top of the sum of the b = 0 s/mm^2^ volumes. Note how each algorithm identifies different voxels, while automatically ignoring voxels belonging to the data or contaminated with signal leakage from multiband imaging.

**Figure 11:**
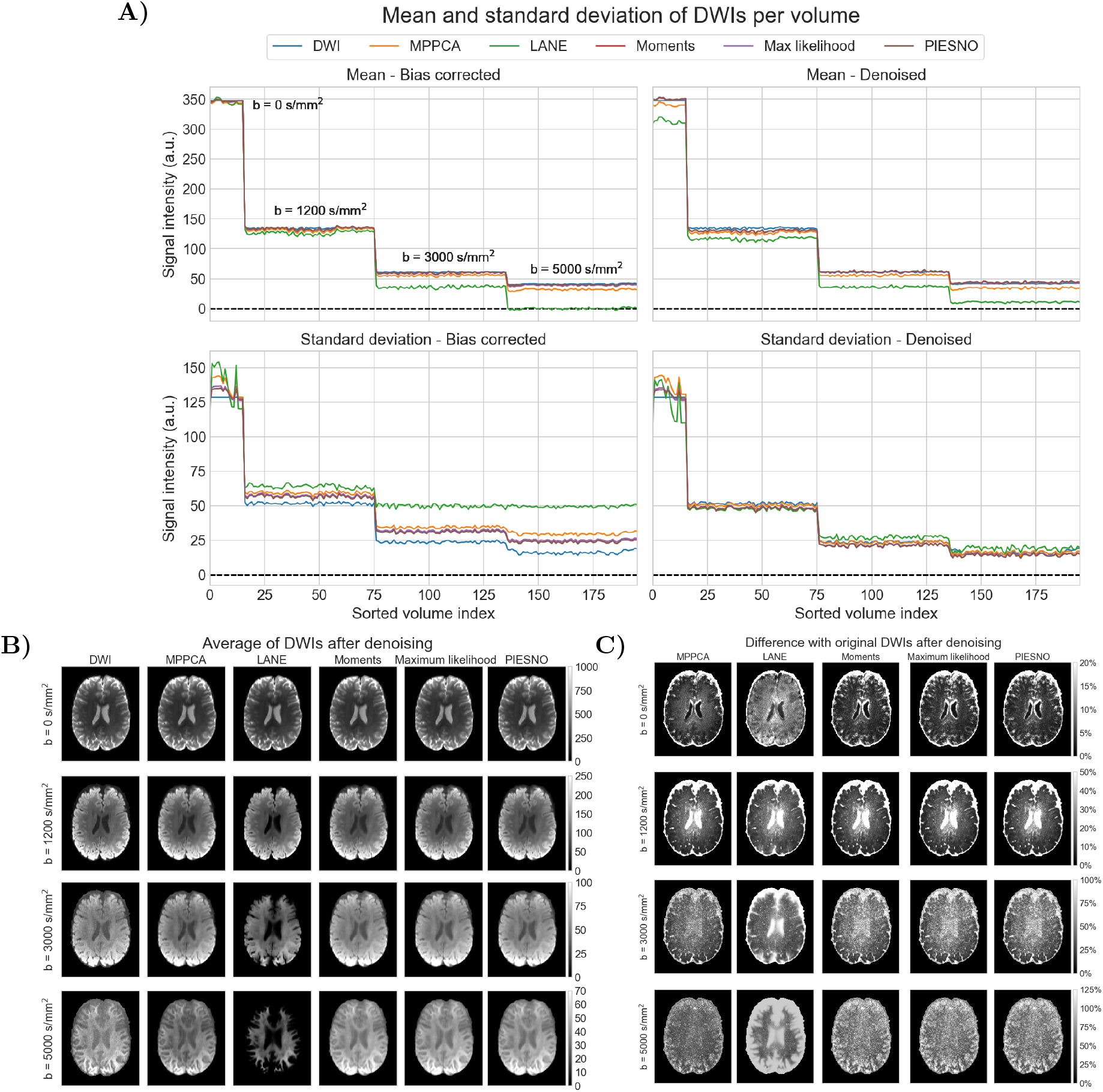
Bias correction and denoising with the NLSAM algorithm of the Connectom dataset from the noise distributions estimated by each method. In **A)**, the left column (resp. right column) shows the result of noncentral chi bias correction (resp. denoising) on the signal value. The top row (resp. bottom row) shows the mean (resp. standard deviation) of the signal inside a white and gray matter mask for each volume. Note how the bias corrected value of LANE goes below 0 (dashed line) due to its high estimation of *σ*_*g*_. After denoising, the standard deviation of the signal decreases as the b-value increases, an effect which is less noticeable for the bias corrected signal only. However, this effect is less pronounced for the bias corrected signal only in the case of LANE and MPPCA. In **B)**, spatial maps of the original data and after denoising (in each column) when averaging datasets at the same b-value for b = 0 s/mm^2^, b = 1200 s/mm^2^, b = 3000 s/mm^2^ and b = 5000 s/mm^2^ (in each row) for each method. Note how each b-value uses a different scale to enhance visualization even though the signal intensity is lower for increasing b-values. Panel **C)** shows the difference in percentage between the original data and after denoising using parameters as estimated by each algorithm.

## 5. Discussion

We have shown how a change of variable to a Gamma distribution *Gamma*(*N*, 1) can be used to robustly and automatically identify voxels belonging only to the noise distribution. At each iteration, the moments (Eqs. (8) and (9)) and maximum likelihood equations (Eqs. (11) and (12)) of the Gamma distribution can be used to compute the number of degrees of freedom *N* and the Gaussian noise standard deviation *σ_g_* relating to the original noise distribution. Voxels not adhering to the distribution are discarded, therefore refining the estimated parameters until convergence. One of the advantage of our proposed methods is that no *a priori* knowledge is needed from the acquisition or the reconstruction process itself, which is usually not stored or hard to obtain in a clinical setting. Results from Section 4.1 show that we can reliably estimate parameters from the magnitude data itself in the case of stationary distributions. For spatially varying distributions without parallel acceleration, the proposed methods achieve an average percentage error of approximately 10% when estimating *σ_g_*, which is equal or better than the other methods compared in this work. Estimated values of *N* are around the true values, even when *σ_g_* is misestimated. While these experiments may still be considered to be simplistic when compared to modern scanning protocols where parallel acceleration is ubiquitous, they highlight that even textbook cases can lead to misestimation if the correct signal distribution is not taken into account. Practical tasks taking advantage of the signal distribution such as bias correction (Pieciak et al., 2018), noise floor removal (Sakaie and Lowe, 2017), deep learning reconstruction with various signal distributions (Lønning et al., 2019) or diffusion model estimation (Collier et al., 2018; Zhang et al., 2012; Landman et al., 2007) may be tolerant, but not perform optimally, to some misestimation of the noise distribution. See e.g., Hutchinson et al. (2017) for discussions on the impact of noise bias correction on diffusion metrics in an ex vivo rat brain dataset. Note that for some applications such as denoising, only the product of the parameters of the distribution might be needed (i.e., 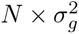) (Pieciak et al., 2016), which is a case we did not cover in the present manuscript.

### Effects of misspecification of the noise distribution

Experiments with SENSE from Section 4.1 reveal that using a local estimation with noise maps provides the best estimates for the proposed methods and PIESNO. MPPCA and LANE perform better when using DWIs as the input rather than noise maps, but at the cost of a broader range of estimated values for *σ_g_* and still underperform when compared to the three other methods. This is presumably because the signal diverges from a Gaussian distribution at low SNR (Gudbjartsson and Patz, 1995) and especially in noise maps, leading to a misspecification of its parameters when the assumed noise distribution is incorrect. Phantom experiments carried with GRAPPA show similar trends except for PIESNO, which overestimates *σ_g_* as shown in Fig. 2. When erroneously fixing *N* = 1, low intensity voxels where *η* > 0 (e.g., gray matter) may be mistakenly included in the distribution after the change of variables, leading to overestimation of *σ_g_*.

The presence of tissue in voxels used for noise estimation might compromise the accuracy of the estimated distributions as shown in Section 4.1. This can be explained by the lower number of noise only voxels available to the proposed methods and PIESNO and to difficulty in separating the signal from the noise for MPPCA and LANE at low SNR. Using measured noise maps is not a foolproof solution as by definition they set *η* = 0, while the (unknown) noiseless signal from tissues is *η* > 0. As the noise distribution may depend on *η* (Aja-Fernández et al., 2014), this means that its parameters (e.g., from a GRAPPA reconstruction) will be inherently different than the one estimated from noise maps. This effect can be seen in Fig. 6, where the estimated values of *N* from noise maps and DWIs are close to 1 for SENSE as expected in theory. For GRAPPA, they are either overestimated and underestimated in regions of the phantom and overestimated in background regions as *N* locally depends on *η*. Accurate estimation of *σ_g_* and *N* over signal regions still remains an open challenge. Nevertheless, the median of the estimated distribution of *σ_g_* is closer to the true distribution when using noise maps than when using DWIs for the proposed methods. Such noise map measurements could therefore provide improved signal distribution estimation for, e.g., body or cardiac imaging, where no intrinsic background measurements are available.

### Effects of parallel imaging and multiband in a phantom

Section 4.2 presented results from a scanned phantom using SENSE coupled with multiband imaging. While no ground truth is available, a SENSE acceleration should provide a Rician signal distribution (*N* = 1) and *σ_g_* should increase with 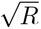 (Aja-Fernández et al., 2014). Fig. 7 shows that for a common SENSE factor, all values of *σ_g_* estimated with *MB* = 3 are larger than at lower factors. The use of multiband imaging should not influence the estimation of *σ_g_* as it only reduces the measured signal, and not the noise component unlike SENSE does. Indeed, estimated values of *σ_g_* are stable until *MB* = 3 or *R* = 2 and *MB* = 2 is used; this is possibly due to signal leakage and aliasing signal from multiband folding over from adjacent slices with higher factors (Todd et al., 2016; Barth et al., 2016). Noise maps are less affected by this artifact, which is already present when *R* = 2 and *MB* = 2, as adjacent voxels have low values, similarly to unaffected voxels. However, leaking signal in DWIs might impact parameters estimation as it can be interpreted as an increase in SNR and therefore a lower noise contribution than expected. Estimation of *σ_g_* is also increasing approximately with 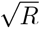 for all methods as expected (Aja-Fernández et al., 2014). While we can not quantify these results, this follows the synthetic experiments with SENSE shown in Fig. 4, where PIESNO and the proposed methods were more precise in estimating *σ_g_* from noise maps.

In the case of estimation using DWIs as input, this expected increase in *σ_g_* for increasing SENSE factor is less obvious e.g., LANE estimates of *σ_g_* decrease from *R* = 2 to *R* = 1 for the no multiband case. As MPPCA and LANE also estimate *η*, it could explain the larger variance of *σ_g_* as *η* fundamentally depends on the microstructural content of each voxel, which is complex and subject to large spatial variations, e.g., notably across DWIs. This also means that estimation over DWIs is susceptible to signal leakage, which would explain the increased estimated values of *σ_g_* for *MB* = 2 and *MB* = 3 for a given SENSE factor. In the noise maps, we have observed that MPPCA and LANE estimated *η* > *σ*_*g*_ in all cases (results not shown). Overestimating the true value of *η* = 0, which is an implicit assumption in PIESNO and the proposed methods, could explain underestimation of *σ*_*g*_ when using noise maps. This overestimation of *η* in turn leads to lower estimates of *σ*_*g*_. The use of multiband and the inherent signal leakage at high factors could explain this overestimation of *η* and underestimation of *σ_g_* for all tested cases. In the case of SENSE, the proposed methods estimated approximately *N* = 1 in all cases, suggesting robustness to multiband artifacts.

### Estimation of noise distributions for in vivo datasets

To complement earlier sections, two datasets acquired on different scanners combining parallel and multiband imaging were analyzed in Section 4.3. Fig. 8 shows that assuming a Rician distribution with *N* = 1 can prove inadequate in some situations. The four repetitions of a single subject acquired on a GE scanner point towards a half Gaussian distribution instead as evidenced by the computed values of *N* around 0.5. This is further evidenced by the low number of voxels (less than 10) detected by PIESNO while assuming *N* = 1. In the preliminary results of our MICCAI submission (St-Jean et al., 2018b), using *N* = 0.5 for PIESNO gave similar results to the proposed methods, suggesting the departure of the data from a pure Rician distribution. Additionally, Fig. 9 shows that those voxels identified automatically as pure noise also adhere closer to a chi distribution than a Rayleigh distribution (where *η* = 0 in both cases). Considering the whole distribution of the data, which is contaminated by artifacts, would also lead to a different distribution. Even if local methods can consider spatially varying noise profile, the local estimation of *σ_g_* will invariably be affected whenever those same artifacts repeat over the data. This introduce a compromise between avoiding artifacts at the cost of reduced spatial specificity and local methods which may not be able to exclude artifacts, but provide local estimations of *σ_g_*. Measurements from noise maps, if available, could therefore offer a middle ground if *N* is low or does not depend locally on the coil geometries (e.g., SENSE or homodyne reconstruction) as shown in Section 4.1.

Fig. 10 shows a large range of estimates for *σ_g_* across methods. In particular, the moments and maximum likelihood equations estimate smaller values of *σ_g_* than MPPCA and LANE, but larger than PIESNO, while still recovering values of *N* close to 1 and successfully discard voxels contaminated by multiband artifacts. The correct value of *σ_g_* most likely sits between these two results as parallel MRI produces spatially varying noise profile, which is higher in the center and not fully captured by the background signal, but the local estimation methods also overestimated *σ_g_* in our synthetic simulations. In panel **A)**, MPPCA and LANE estimates of *σ_g_* with DWIs are likely affected by multiband artifacts as the median is larger than the signal level at b = 5000 s/mm^2^. This indicates a possible overestimation as *σ_g_* should be lower than the measured signal at the highest b-value. For PIESNO and the proposed methods, the median *σ_g_* is lower than the median of the reference b = 5000 s/mm^2^ data. An overestimation of *N* could explain the low values of *σ_g_* estimated by the proposed methods just as misestimation of *η* by MPPCA and LANE could affect their respective estimate of *σ_g_* by balancing out the misestimated values.

Fig. 11 shows the result of each method on a bias correction and denoising task on the Connectom dataset. In panel **A)**, the standard deviation of the signal (bottom left panel) is increased after bias correction for LANE (green line) and decreased (around the same level) for the other methods when compared to the uncorrected data (blue line). The situation is similar after denoising, but to a lesser extent, while the moments, maximum likelihood equations and PIESNO follow the same signal level as the unprocessed data on average. Regarding the mean of the signal itself, LANE is on average lower or close to 0 after bias correction, indicating potential degeneracies due to overestimation of *σ_g_*. From panels **B)** and **C)**, the results of all methods are visually similar except for LANE (especially at b = 3000 s/mm^2^ and b = 5000 s/mm^2^), indicating that the NLSAM denoising algorithm treated different values of *σ_g_* in the same way. This is because the optimal regularized solution (which depends on *σ_g_*) is piecewise constant (St-Jean et al., 2016; Tibshirani and Taylor, 2011) and can tolerate small deviations in *σ_g_*. Finally, MPPCA, the moments and maximum likelihood equations and PIESNO perform similarly, even if they estimated different values of *σ_g_* and *N*, with MPPCA showing slightly lower signal intensity at b = 5000 s/mm^2^. This could be due to the bias correction having a larger effect when *σ_g_* is larger, increasing the standard deviation of the resulting signal. As shown in panel **C)**, the difference with the original dataset for MPPCA is lower than the proposed methods or PIESNO, even though the estimated value of *σ_g_* was larger.

## 6. Conclusions

We presented a new, fully automated framework for characterizing the noise distribution from a diffusion MRI dataset using the moments or maximum likelihood equations of the Gamma distribution. The estimated parameters can be subsequently used for e.g., bias correction and denoising as we have shown or diffusion models taking advantage of this information. This requires only magnitude data, without the use of dedicated maps or parameters intrinsic to the reconstruction process, which may be challenging to obtain in practice. The proposed framework is fast and robust to artifacts as voxels not adhering to the noise distribution can be automatically discarded using an outlier rejection step. This makes the proposed methods also applicable on previously acquired datasets, which may not carry the necessary information required by more advanced estimation methods. Experiments using parallel MRI and multiband imaging on simulations, an acquired phantom and in vivo datasets have shown how modern acquisition techniques complicate estimation of the signal distribution due to artifacts at high acceleration factor. This issue can be alleviated with the use of noise only measurements or by limiting the acceleration factor to prevent signal leakage. Moreover, different vendors implement different default reconstruction algorithms which leads to different signal distributions, challenging the strategy of assuming a Rician distribution or approximations of *N* based on the physical amount of channels in the receiver coil. We also have shown how signal bias correction and denoising can tolerate some misestimation of the noise distribution using an in vivo dataset. Noteworthy is that the theory we presented also applies to any other MRI weighting using large samples of magnitude data (e.g., functional MRI, dynamic contrast enhanced MRI). This could help multicenter studies or data sharing initiatives to include knowledge of the noise distribution in their analysis in a fully automated way to better account for inter-scanner effects.

## Appendix A. Estimating parameters of the Gamma distribution

## Estimation using the method of moments.

For any given distribution, we can estimate its parameters by relating the samples and the theoretical expression of its moments. The Gamma distribution is parametrized as *Gamma*(*α, β*) and has a probability distribution function of

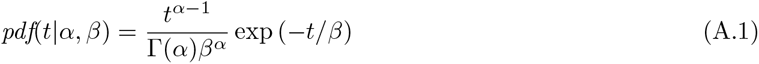

with *t, α, β* > 0 and Γ(*x*) the gamma function. The first moments are analytically given by (Chap. 5 Papoulis, 1991; Weisstein, 2017).

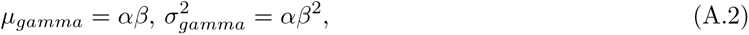

In this paper, the Gamma distribution parameters are *Gamma*(*α* = *N, β* = 1) after the change of variable 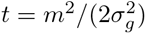 for our particular case. Since we have *β* = 1, this leads to a special case where the mean and variance are *equal* with a value of *α* = *N* and can be expressed only in terms of the magnitude signal *m*. For simplicity, we will only use the mean *µ_gamma_* and variance 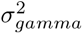 to estimate the required parameters *N* and 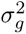, but higher order moments could also be used. However, in practice, they might accumulate numerical errors due to the higher powers involved and are not used here since two equations are enough to estimate the two parameters. Starting from the analytical expression given by Eq. (A.2), we have for the special case *Gamma*(*N*, 1)

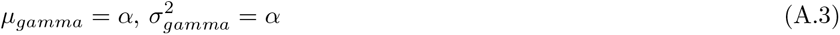

Which we can compute using the sample mean and sample variance formulas such that

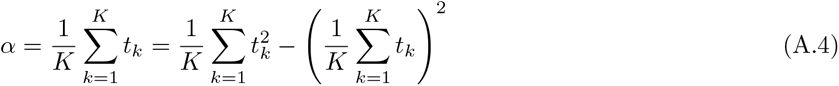

Substituting the equation for the moments in terms of 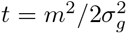, we obtain

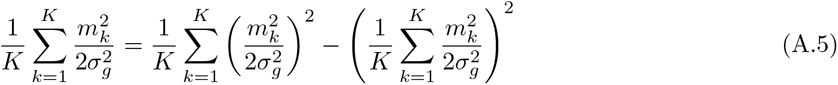

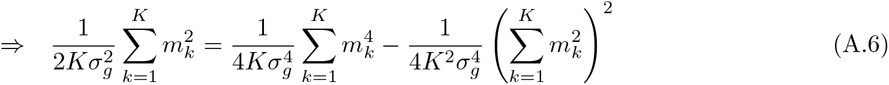

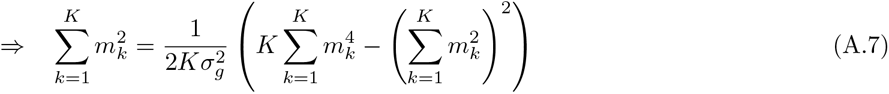

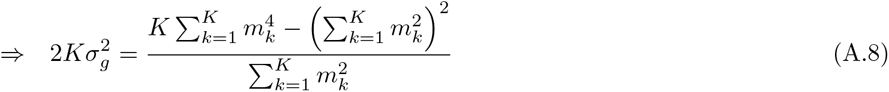

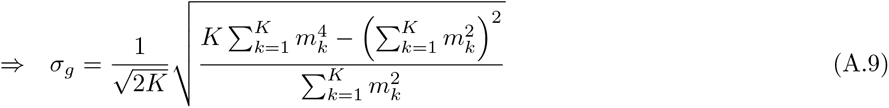

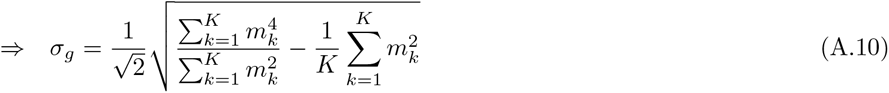

Therefore, it is possible to estimate the Gaussian noise standard deviation using Eq. (A.10) and the values of magnitude data *m_k_*, assuming that the voxels considered here do not contain any object signal. With the value of the noise variance 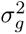 now known, going back to the original Gamma distribution *Gamma*(*α* = *N, β* = 1) yields the number of coils *N* as previously shown by Eq. (9)

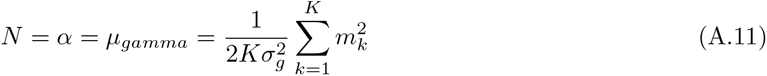

## Estimation using maximum likelihood equations

An alternative to the method of moments to estimate parameters from a given distribution is to solve the equations derived from its likelihood function for each unknown parameter. Given a set of observed data, maximizing the likelihood function from a known distribution (or equivalently, the log of the likelihood function) yields a set of equations to estimate its parameters. For the *Gamma*(*α, β*) distribution, maximizing the log likelihood by equating the partial derivative to 0 for each parameter yields (Thom, 1958)

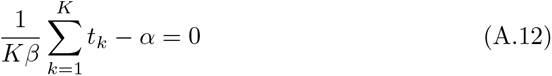

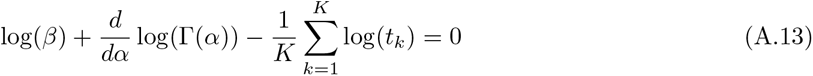

Since we have *α* = *N* and *β* = 1, in this special case Eq. (A.12) is the same as Eq. (A.11).

Combining Eqs. (A.12) and (A.13) yields an implicit equation to estimate *σ_g_*, which can be written as

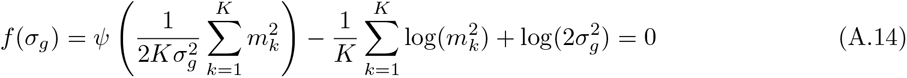

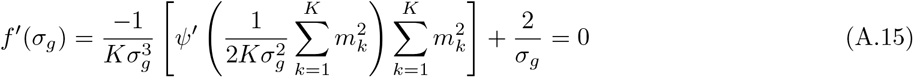

and Eq. (A.13) can be rewritten as an implicit equation of *N*

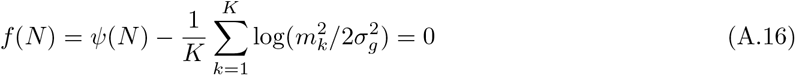

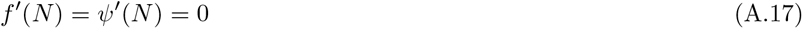

where 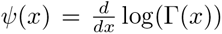 is digamma function and *ψ*′ is the derivative of *ψ*, called the polygamma function. Eqs. (A.15) and (A.17) can be solved numerically using Newton’s method provided we have a starting estimate *x*_0_. The update rule for Newton’s method at iteration *n* is therefore

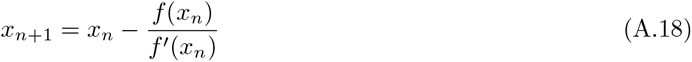

For the first iteration, a starting estimate *x*_0_ to approximate the solution is needed. For Eq. (A.15), we use *x*_0_ = *σ_m_*, where *σ_m_* is the sample standard deviation of the identified noise only voxels. A starting estimate for Eq. (A.17) is given by (Minka, 2012) considering 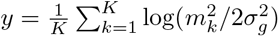.

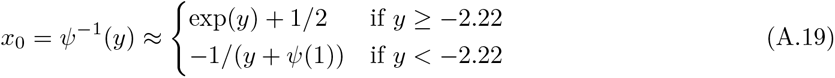

In practice, we have observed that 5 iterations of Eq. (A.18) were sufficient to reach |*x*_*n*_ − *x*_*n*−1_| < 10^−13^.

## Appendix B. Generalized bias correction

As an application which requires knowledge of both *σ_g_* and *N*, we now present a general version for non integer values of *N* of the signal bias correction from Koay and Basser (2006); Koay et al. (2009a). The correction factor *ξ*(*η*|*σ*_*g*_, *N*) can be used to obtain *η* from the magnitude measurement *m*_*N*_ given the values of *σ*_*g*_ and *N* such that

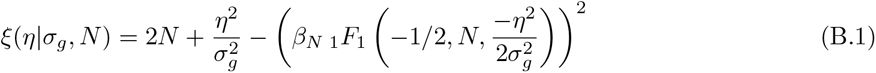

where _1_*F*_1_ is Kummer’s function of the first kind. By defining

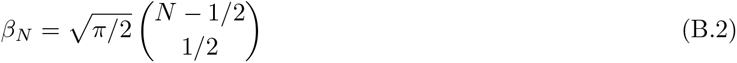

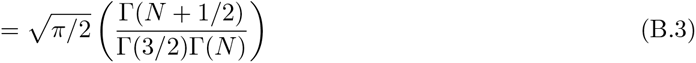

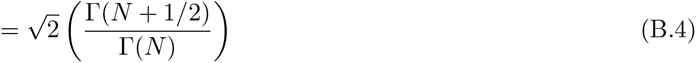

where 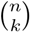 is a binomial coefficient, we obtain a generalized version of Eq. (B.1) which can now be applied for non integer values of *N*, such as in the case of a half Gaussian signal distribution (*N* = 0.5) which occurs when employing half-Fourier reconstruction techniques (Dietrich et al., 2008). Estimation of *η* is finally done with

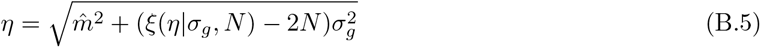

where 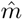 is an estimate of the first moment of a noncentral chi variable and is estimated from a spherical harmonics fit of order 6 on the DWI datasets for each shell in the present work. Eq. (B.5) can be solved iteratively w.r.t. *η* until convergence, see (Koay et al., 2009a) for further implementation details.

## Appendix C. Automated identification of noise only voxels

This appendix outlines the proposed algorithm and details for a practical implementation. Our implementation is also freely available at https://github.com/samuelstjean/autodmri (St-Jean et al., 2019) and will be a part of ExploreDTI (Leemans et al., 2009). The synthetic and acquired datasets used in this manuscript are also available (St-Jean et al., 2018a).

**Algorithm 1:**
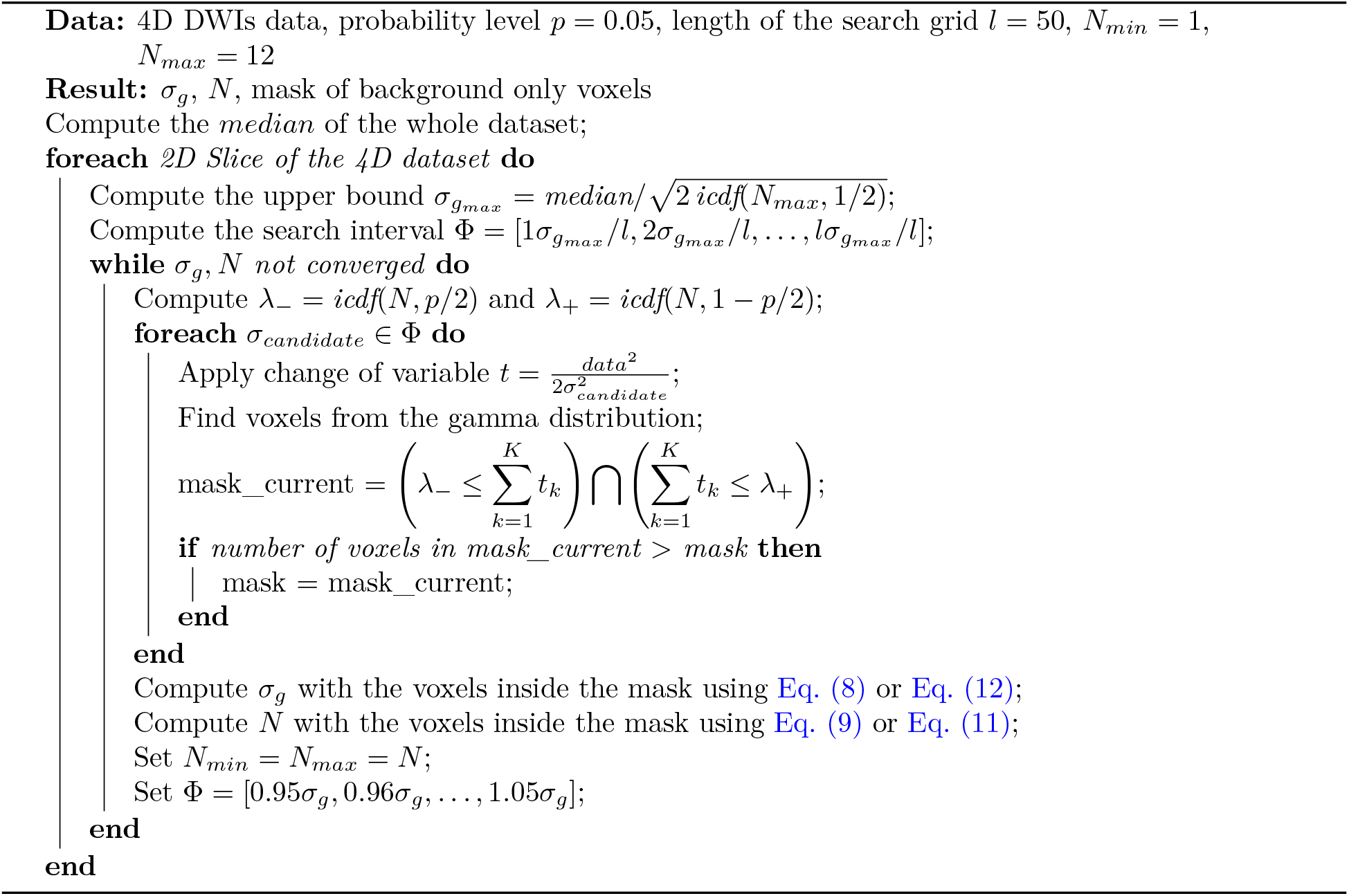
Main algorithm to identify voxels belonging to the Gamma distribution

## Acknowledgements

We would like to thank Michael Paquette for useful comments and discussion. The authors have declared no conflict of interest. The funding agencies were not involved in the design, data collection nor interpretation of this study. This research is supported by the Netherlands Organization for Scientific Research (NWO), Grant/Award Number: VIDI 639.072.411. Samuel St-Jean is supported by the Fonds de recherche du Québec - Nature et technologies (FRQNT) (Dossier 192865) and Chantal M. W. Tax is supported by the Netherlands Organization for Scientific Research (NWO), Grant/Award Number: Rubicon 680-50-1527.

## Footnotes

1 The inverse cdf is also known as the quantile function.

2 As there is no analytical solution to the inverse cdf of a Gamma distribution, one can use the function gaminv(*p, α, β* = 1) in Matlab or InverseGammaRegularized(*α*, 1 − *p*) in Mathematica to numerically estimate it.

3 https://github.com/samuelstjean/autodmri

4 https://mathworks.com/matlabcentral/fileexchange/36893-parallel-mri-noisy-phantom-simulator

5 https://openfmri.org/dataset/ds000031

6 https://www.cardiff.ac.uk/cardiff-university-brain-research-imaging-centre/research/projects/cross-scanner-and-cross-protocol-diffusion-MRI-data-harmonisation

## Notes

#### Summary of Updates

v3: Peer reviewed version v2: Revised for submission to Medical image analysis v1: Initial submission to Magnetic resonance in medicine

https://zenodo.org/record/2483105

https://github.com/samuelstjean/autodmri

