## Supplementary materials for "Automated characterization of noise distributions in diffusion MRI data"

Table S1 presents the URL where to find the software packages and the version used in this manuscript. Unless indicated in the manuscript, default parameters set by the respective software were used. Experiments were carried on a standard desktop computer with an Intel i7 processor running Ubuntu Linux 16.04.

Note also that the raw data we generated and the scanned phantom datasets used in this manuscript are also available online at http://dx.doi.org/10.5281/zenodo.2483105

| Algorithm | URL | Version | DOI for the implementation |
|---|---|---|---|
| Bias correction | https://github.com/samuelstjean/nlsam | v0.6.1 | |
| LANE | https://cran.r-project.org/web/packages/dti/index.html | 1.2 | |
| Maximum likelihood | https://github.com/samuelstjean/autodmri | v0.1 | 10.5281/zenodo.3339157 |
| Moments | https://github.com/samuelstjean/autodmri | v0.1 | 10.5281/zenodo.3339157 |
| MPPCA | https://github.com/MRtrix3/mrtrix3 | 0.3_RC1 | |
| NLSAM | https://github.com/samuelstjean/nlsam | v0.6.1 | |
| PIESNO | https://github.com/dipy/dipy | 0.12 | |

Table S1: Software version used in this manuscript. We additionally provide DOIs to the source code for our own implementation as it leads to a curated archive of the latest available version automatically.

*Corresponding author
